# Viral impacts on microbial activity and biogeochemical cycling in a seasonally anoxic freshwater lake

**DOI:** 10.1101/2023.04.19.537559

**Authors:** Patricia Q. Tran, Samantha C. Bachand, Benjamin Peterson, Shaomei He, Katherine McMahon, Karthik Anantharaman

## Abstract

Microbial biogeochemical cycling relies on alternative electron acceptors when oxygen is unavailable, yet the role of viruses (bacteriophages) in these processes is understudied. We investigated how seasonal anoxia impacts viral and microbial biogeochemical cycling, by using paired total metagenomes, viromes, and metatranscriptomes, that were collected weekly. Stratification and anoxia drove microbial community composition, but dataset origin impacted the interpretation of viral community structure, activity, and function. Importantly, taxa abundance did not correlate with activity for both microbes and viruses. We identified virus-host linkages for 116 phages across 55 distinct hosts, many of which expressed genes for aerobic methane oxidation, nitrogen fixation, denitrification, and sulfate reduction. Overall, this work demonstrates the breadth and dynamics of virus-host interactions in mediating biogeochemistry. Additionally, we propose that viral community detection, functional potential, and activity are sensitive to pre-sequencing decisions, which must be kept in mind when interpreting genomic data in a biologically meaningful way.

## Introduction

Anoxic zones are globally distributed and span oceans, coastal zones, and lakes. Lakes are expected to become more anoxic in coming decades, as warming water surface temperatures will increase temperature/density differences between the top and bottom layers of lakes, making it harder for water to mix throughout^1–3^. The increase in the duration and extent of anoxic zones is expected to transform biogeochemical cycling. Microbes play critical roles in carbon, nitrogen, sulfur, and phosphorus cycling under anoxia^4^. For example, under anoxic conditions, microbes transform inorganic mercury to toxic methylmercury^5, 6^, reduce sulfate to toxic hydrogen sulfide, reduce nitrate by denitrification to form the potent greenhouse gas, nitrous oxide, and conduct methanogenesis to form the greenhouse gas, methane.

In seasonally anoxic lakes, the upper mixed water column is a habitat for fish, photosynthetic organisms, and aerobic heterotrophic microbes. In contrast, anoxic habitats are home to microbes with more diverse metabolic strategies such as chemolithotrophy, which is the use of inorganic compounds to generate energy and synthesize organic carbon. In addition, in the absence (or under the lower pressure) of top-down control of grazers and eukaryotic bacterivores on the bacterial community in anoxic hypolimnion, viral influences in controlling microbial communities can be heightened^7^. When considering the whole microbial community (viruses, bacteria, archaea, and eukaryotes), culture-independent metagenomics has contributed to the discovery and inference of virus-host relationships. Metagenomics can hint to which processes might be happening in the environment, with the caveat that genes must be transcribed into RNA, modified based on the cellular environment, and form proteins. For example, cyanophages in marine habitats have been shown to control photosynthesis^8^. The use of viral genomics also resulted in insights about the viral communities of freshwater lakes^9–13^. These approaches can be supplemented by (meta)transcriptomics (RNA-seq) and cultivation-based studies to test these predicted viral functions in the environment.

Besides the need for further lines of evidence to support metagenomic data (e.g., transcriptomics, field-based measurements, or physiology studies), is the need for benchmarking viral omics methods. Two major methodologies are commonly applied to identify viral genomes depending on where/how the viral fraction is obtained: 1) bioinformatically mining “total” metagenomes (microbial cells plus viruses associated with cells) to extract viral genomes (presumably recovering viruses attached to or inside host cells, also referred to as cellular metagenomes in this study), and 2) physically collecting and sequencing the cell-free fraction (presumably largely consisting of free viral-like particles, referred to as viromes in this study). Each method has its own theoretical biases, but the extent to which these biases can affect the interpretation of viral-host interactions in freshwater lakes is unclear because previously published freshwater studies mainly rely on bioinformatically extracting viral genomes from the total metagenomic fraction. Computationally extracting viruses from total metagenomes is a common way to “discover” viruses and make ecological interpretations. However, it is unknown how this methodological choice influences ecological interpretation specifically in freshwater environments. For example, in agricultural soils, viromes outperform total metagenomes in assessing the viral community, but this conclusion may or may not be applied to freshwaters, as environmental factors such as moisture, pH, availability of organic matter or recalcitrant material may affect DNA extraction may be specific to each environment^14^. Given that total metagenomes are often used to identify viral metagenomic scaffolds and to annotate functions (e.g., auxiliary metabolic genes (AMGs)), it is important to understand how methodology choice affects ecological interpretations (e.g., AMGs involved in photosynthesis and biogeochemical cycling of nitrogen, sulfur, and carbon).

Lake Mendota (Madison, Wisconsin, USA) is a eutrophic freshwater lake that has a long-term continuous monitoring program focused on microbial communities in the upper mixed layer^15, 16^ and some studies of the microbial community in the anoxic hypolimnion (deep waters)^6^. Prior work shows that the microbial community in hypolimnion is different compared to the epilimnion, and has distinct biogeochemical functions^6, 17^. Here, we seek to understand how environmental heterogeneity (defined as spatiotemporal patterns of anoxia, and pre-post fall mixing) impacted virus-host interactions, and how these interactions in turn impacted their activity in carbon, sulfur, and nitrogen cycling.

## Results

We collected depth-discrete samples from Lake Mendota between July 16, 2020, to October 19, 2020, spanning the period of thermal stratification and fall mixing. Lake Mendota was thermally stratified from late May until October 18, 2020 (**Figure 1A**). Anoxic water was detected in July, as is historically typical for Lake Mendota^1, 18^. We chose 16 water samples that captured the stratified and mixed period for sequencing (16 total metagenomes, 14 viromes, 16 metatranscriptomes) **Supplementary Table 1**. Samples captured a range of environmental data such as oxygen (including oxic, oxycline, anoxic layers), pH, phycocyanin, turbidity, and water temperature.

**Figure 1.**
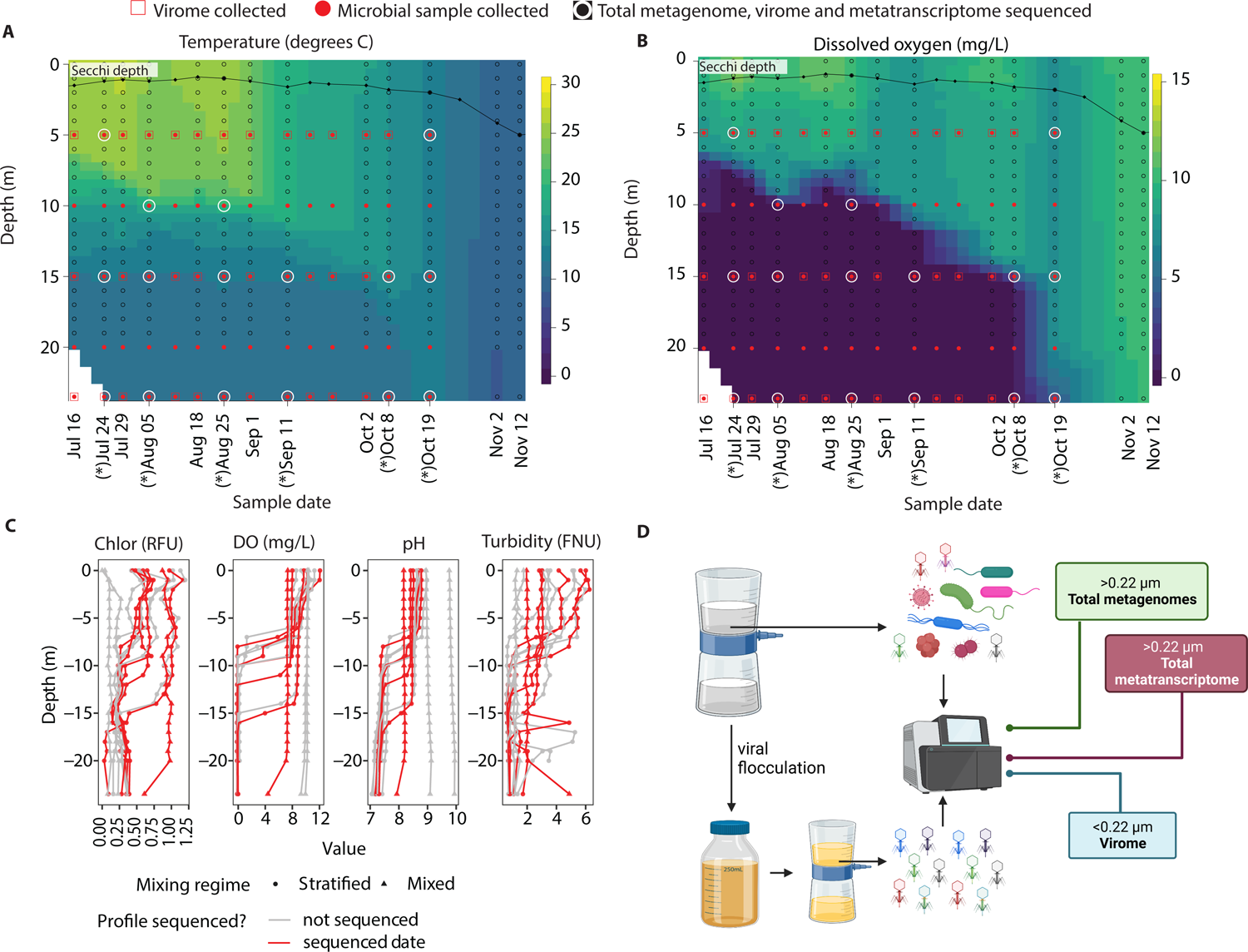
Samples and environmental data. Weekly samples were collected at 5, 10, 15, 20, and 23.5m from July 16, 2020, to October 19, 2020. Viral samples were obtained for depths 5, 15, and 23.5m each week. Samples were collected during the day. **A. and B.** Temperature (A) and dissolved oxygen profiles (B) in Lake Mendota. The lake was stratified until fall mixing on October 18, 2020. 16 samples representing the oxic, anoxic, and oxycline layers were selected for total metagenome, virome, and metatranscriptome sequencing. Asterix represent dates where samples were sequenced. **C.** Chlorophyll (Chlor), dissolved oxygen (DO), pH, and turbidity profile in 2020. **D.** From each sample, water was filtered through 0.22μm filters for total metagenomes and metatranscriptomes and the filtrate was subject to viral precipitation with iron chloride to generate subsequent viromes.

While we did not collect sulfide data in 2020, data collected in 2021 showed that sulfide began accumulating in the water column in mid-summer and accumulated in the anoxic hypolimnion during the stratification period. Lake Mendota typically has sulfide concentrations higher than 150μM in the anoxic hypolimnion (**Supplementary Figure 1**) and substantial oxygen demand due to high epilimnion primary productivity^19^. Lake Mendota is a low-iron and high-sulfate lake (sulfate is of geological origin) and prior studies have documented vigorous sulfate-reduction^20^, based on measurements in the sediment^21^, and from the profile of sulfate-sulfur in the anoxic hypolimnion^6, 17^. While we did not collect methane and nitrate and nitrite in 2020, prior studies have shown the accumulation of methane during summer stratification^22^ and nitrate/nitrite in the water column.

### Microbial community composition and expression change as a function of anoxia, but abundance does not correlate to activity

We assembled metagenome-assembled genomes (MAGs) from the 16 total metagenomes and identified viral genomes in both the total metagenome and virome fractions, and used bioinformatic analyses to link phages to their potential bacterial hosts. Finally, we used metatranscriptomes to assess the activity of these phage-host pairs. We obtained a final set of 431 MAGs (429 bacteria, 2 archaea) of medium to high quality for downstream analyses. Among them, 349 MAGs were assigned to 176 unique genera and 82 MAGs could not be assigned to any genera (**Supplementary Table 2, Supplementary Figure 2**). Of those 82 MAGs without a genus assignment, most of them were associated with the phyla Verrucomicrobiota, Bacteroidota, and Patescibacteria. 133 MAGs shared ≥ 97% ANI (average nucleotide identity) to MAGs previously published from Lake Mendota^4, 6^. Notably, previous MAGs were generated from different water samples, namely water column integrated from the surface down to 12m^4^, and two depth-discrete anoxic profiles collected in 2017^6^ (**Supplementary Table 3**). Besides common freshwater phyla, we recovered 18 Patescibacteria, 81 Gammaproteobacteria, 42 Kirimatiellaeota, and 20 Verrucomicrobiota MAGs.

The hypolimnion harbored more diverse taxa over time during the stratified period, as indicated by the increase in the Shannon diversity indices (**Figure 2A**). Among site categories, beta-diversity was 0.16 in the oxycline & stratified sample, 0.18 in the anoxic & stratified sample, and 0.20 in the oxic & mixed sample.

**Figure 2.**
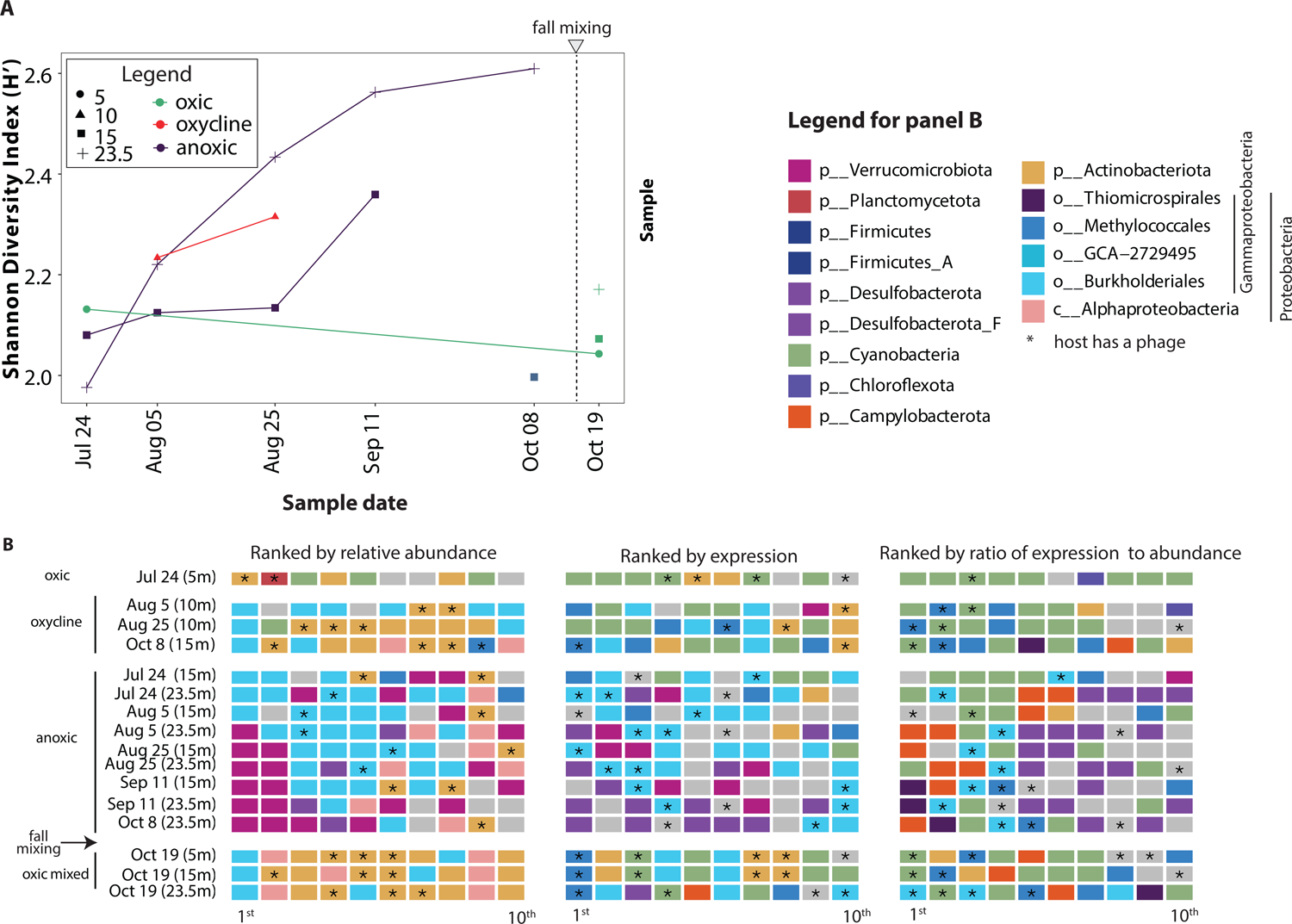
Diversity and the top 10 MAGs ranked by abundance, expression, and overexpression. **A.** The Shannon Diversity indices were calculated in each of the 16 total metagenomes. **B.** Top 10 MAGs were ranked by the abundance (read coverage within a metagenome), expression (RPKM divided by internal standard count), and overexpression (the ratio of expression to abundance), respectively, and were colored by their taxonomy (with Grey indicating taxa not listed in the legend). MAGs with at least one associated phage (**Figure 7**) are indicated with an asterisk.

To determine which organisms were active, we compared MAG abundances to their transcriptional activity. We used the MAG relative abundance, expression, and the ratio of expression to abundance to compare who is present, active, and overexpressed respectively (**Figure 2B**). From metagenome data, we calculated the relative abundance of MAGs within each sample based on their metagenome read coverage, and the top 10 most abundant MAGs in each metagenome are shown in the left panel of **Figure 2B**. From the metatranscriptome data, after mapping, the read counts were first normalized by the internal standard read count (see Methods) to obtain an RPKM_normalized_ value for each MAG, allowing the comparison of expression/activities among different samples. The top 10 most active MAGs in each metatranscriptome are shown in the middle panel of **Figure 2B**. The RPKM_normalized_ value of each MAG was then divided by its relative abundance within the corresponding metagenome, to obtain a ratio of RPKM to abundance to see if a MAG is overexpressed compared to other expressed MAGs. The top 10 overexpressed MAGs are shown in the right panel of **Figure 2B**).

Overall, active taxa were not correlated with abundant taxa. For example, differences in microbial community composition existed among zones of different oxygen availabilities, and the community composition homogenized upon fall mixing as indicated by the similar community composition throughout the water column post-mixing (**Supplementary Figure 3**). In the surface oxygenated samples, the microbial community was dominated by Actinobacteriota, Cyanobacteria, and Planctomycetota, but Cyanobacteria and Actinobacteriota were the most active. At the oxycline, *Burkholderiales* (Gammaproteobacteria) was both abundant and active, although *Methylococcales* (also a Gammaproteobacteria) were among the most active and most overexpressed. In the anoxic hypolimnion, the most abundant taxa were initially *Burkholderiales*, but over time, Verrucomicrobiota became consistently highly ranked. However, Desulfobacteriota was most often highest expressed in the hypolimnion. Additionally, when considering overexpression (high RPKM to abundance ratio), Campylobacterota (f *Sulfurimonadaceae*, g *Sulfuricurvum*) and *Thiomicrospirales* (Gammaproteobacteria) were notable in the anoxic hypolimnion. Both these groups of organisms are known to participate in sulfur cycling and are commonly found in oxygen-minimum zones in marine systems^23–25^. *Sulfuricurvum* is a facultative anaerobic chemolithoautotrophic sulfur oxidizer that has been known to use nitrate as an electron acceptor^26^. *Thiomicrospirales* have been shown to increase with depth in oxygen minimum zones^25^. After fall mixing, the microbial community “homogenized” across layers, with Alphaproteobacteria becoming highly ranked in terms of abundance but in comparison, *Methylococcales* remained a highly expressed taxa, along with some taxa from the hypolimnion (e.g. *Desulfobacteria*). Many of the top-ranked active taxa in the anoxic hypolimnion were associated with phages, as shown by the asterix in **Figure 2B**.

### Viral community composition

The microbial community composition and expression patterns were distinct after fall mixing and differed between the epilimnion and hypolimnion, but the viral community was mostly distinguished by the origin of the dataset analyzed – in other words, whether the viruses were recovered from the total metagenomes or viromes (**Figure 3**). *Caudoviricetes* (*Myoviridae*, *Siphoviridae,* and *Podoviridae*) dominated each sample (**Supplementary Figure 4**). In the viromes, a higher number of viral genomes were of unknown taxonomy, compared to those in the total metagenomes. *Microviridae* contributed to a higher percentage of the total metagenome viral community but were seldom observed in the viromes.

**Figure 3.**
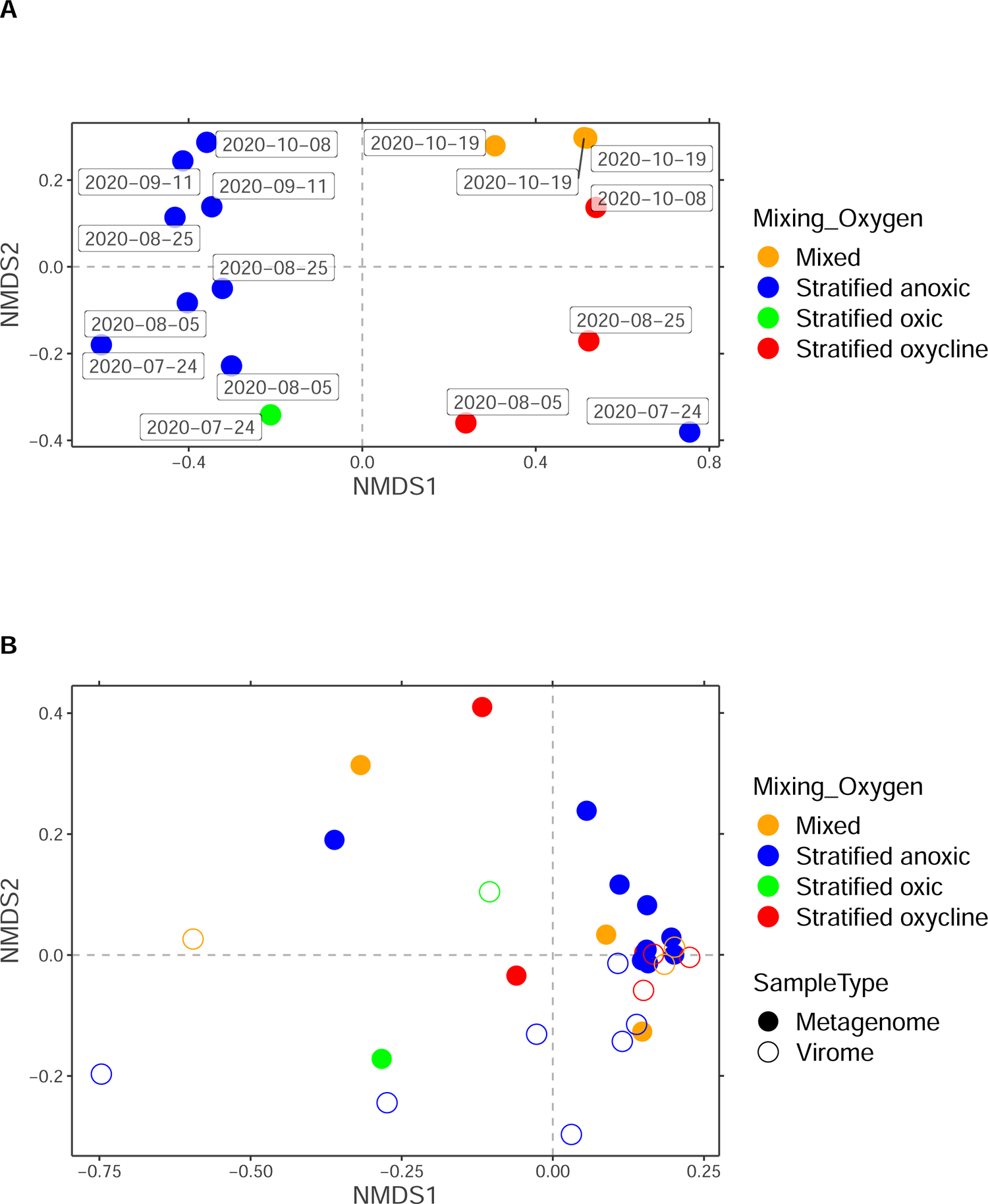
Non-metric dimension scale plot of (A) the prokaryotic community (B) and the viral community. Environmental type (season and stratification) was a driver of the prokaryotic community (clustered by color), but not the viral community. Rather, the viral community was broadly clustered by sample type, that is whether they were originating from a total metagenome or viromes.

### Comparison of viruses in “total metagenomes” vs. “viromes”

Total metagenomes are the fraction that is retained on 0.22 μm filters, and viromes are the fraction that passed through 0.22 μm filters and were chemically precipitated. We investigated the viral community in both the virome and total metagenome fractions to identify differences or similarities in viral community composition, function, and activity.

For any pair of samples, significantly more viral scaffolds were identified in the viromes than in the total metagenomes (paired t-test, t = −11.236, df = 13, p-value = 4.581e-08) (**Figure 4A**). For example, about 60-80% of scaffolds were viral in the viromes, whereas only ∼20% of scaffolds were viral in the total metagenomes. Viromes yielded more phage genomes in most sample pairs, except in 2 sample pairs (August 25^th^ 23.5m and October 10^th^ 23.5m) (**Figure 4B**). Additionally, there tended to be more lytic phages than lysogenic phages in both total metagenomes and viromes (**Figure 4C**).

**Figure 4.**
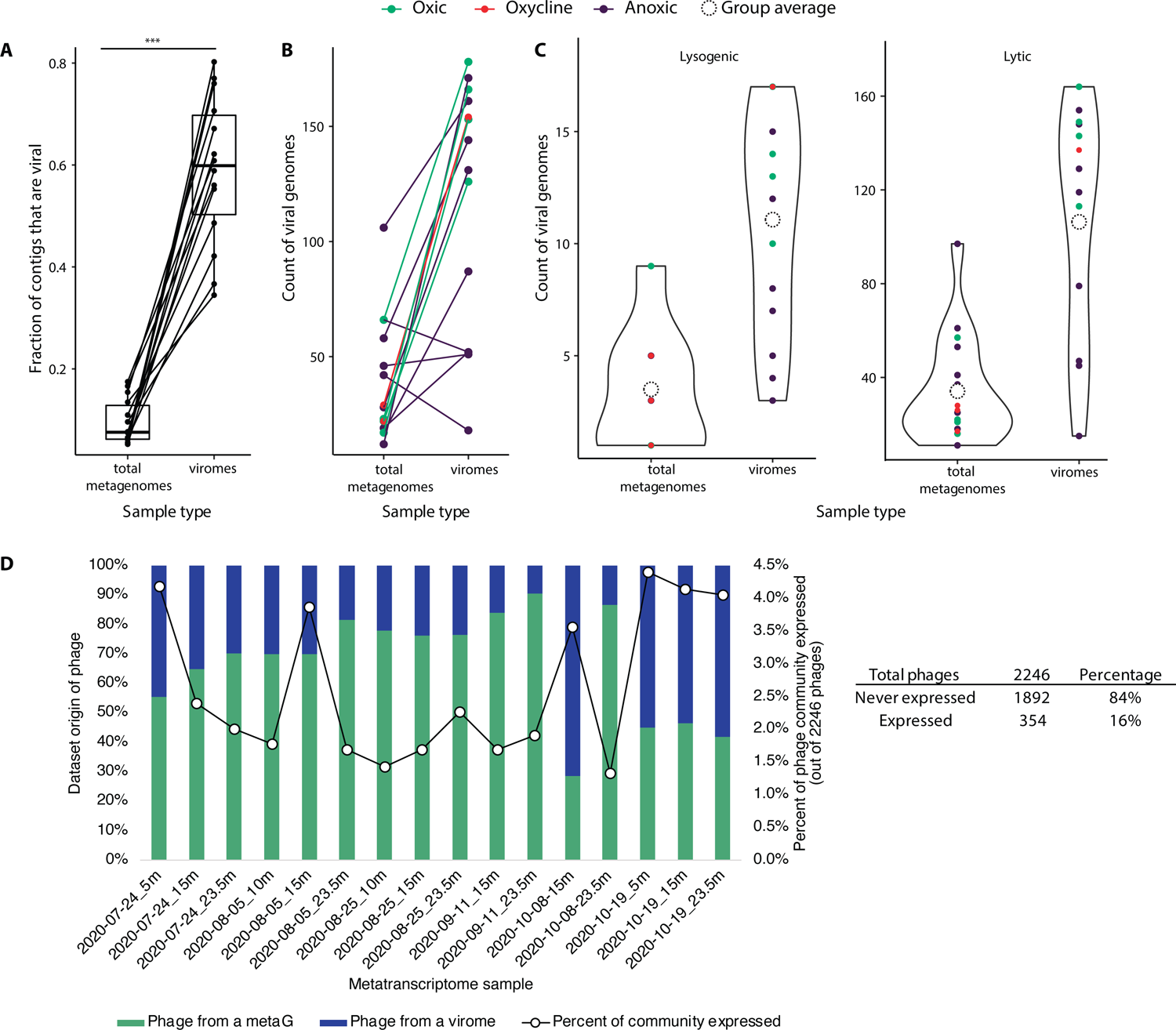
Comparison of viruses based on dataset origin. **A.** A significantly higher percentage (paired t-test, t = −11.236, df = 13, p-value = 4.581e-08) of assembled scaffolds are identified as “viral” from the virome fraction for any given physical sample pair. **B.** The number the putative complete or high-quality phage genomes is higher in the virome than in the total metagenome, except in 2 samples. **C.** The number of lytic and lysogenic phage genomes (complete or high-quality) found in viromes is higher than when computationally extracted from total metagenomes, and the number of recovered lytic phage genomes is higher than the number of lysogenic genomes in both total metagenomes and viromes (note the difference in y-axis scale). **D.** Origin and activity of phages. Of the 2246 phages with complete and high-quality genomes, 16% of them were expressed at least once in the metatranscriptomes. Per sample point, at most ∼4.5% of the 2246 phages were active.

Additionally, we found that the number of complete and high-quality viral genomes was higher in viromes than in the total metagenomes (1,466 vs. 780). The number of complete genomes was higher in total metagenomes than in viromes (176 complete genomes vs. 132). We found 6 mega-phages (>200kpb) and 469 small phages (<15kbp) (all are complete or high-quality genomes) (**Supplementary Table 4**). With 2,246 complete and high-quality phage genomes considered in this analysis, 84% of them were never expressed over the course of the metatranscriptomics time series. 354 (or 16%) of the phages were expressed at least once. On any given day, between 1.34 to 4.41% of the viral genomes showed positive RNA expression (**Figure 4D**). For any given metatranscriptome, a variety of proportions of sample origin existed, but the oxycline (e.g. Oct 08 at 15m) and post-mixing samples tended to have over 50% of active phage originating from the viromes (**Figure 4D**).

### Phage activity and impact on nutrient cycles

Auxiliary metabolic genes (AMGs) are genes found on phage genomes, that are not involved in critical functions like phage replication. While they have different definitions, they are generally considered to augment their hosts’ metabolism, by enhancing certain metabolic pathways that result in more viral progeny being released upon host-cell lysis^27^. 91 different AMG families were identified (distinct KO identifiers) and comprised 1,287 proteins distributed on 494 phage genomes. 135 phages were from identified metagenomes (18 lysogenic, 117 lytic) and 359 were from viromes (28 lysogenic and 331 lytic). We calculated the abundance and RNA expression of those AMG-encoding phages (**Figure 5**). 59 AMGs were found on phages that were inactive (no RPKM expression during any sampling point), and 27 AMGs were on phages that were expressed at least once. Among those that were expressed, 13 AMGs were on phages recovered from the total metagenome fractions, and 6 were on phages recovered only from the virome fractions, and 8 from both fractions. (**Supplementary Table 5**). Notably, photosynthesis AMGs (*psbAD*) were found on phages from both fractions but were transcriptionally inactive. They were identified on phage genomes that could be attributed to 8 closely related species of *Synechococcus* reference hosts. The *pmoC* AMG associated with aerobic methane oxidation was found on an inactivate phage, for which the host could not be identified in our study, but is likely a *Methylococcales* based on a previous study using Lake Mendota data^28^. Organosulfur metabolism AMG (*cysH*) were found and active in both fractions. Finally, a sulfur oxidation AMG (*soxB*) was found in viral genomes originating from the viromes but transcriptionally inactive.

**Figure 5.**
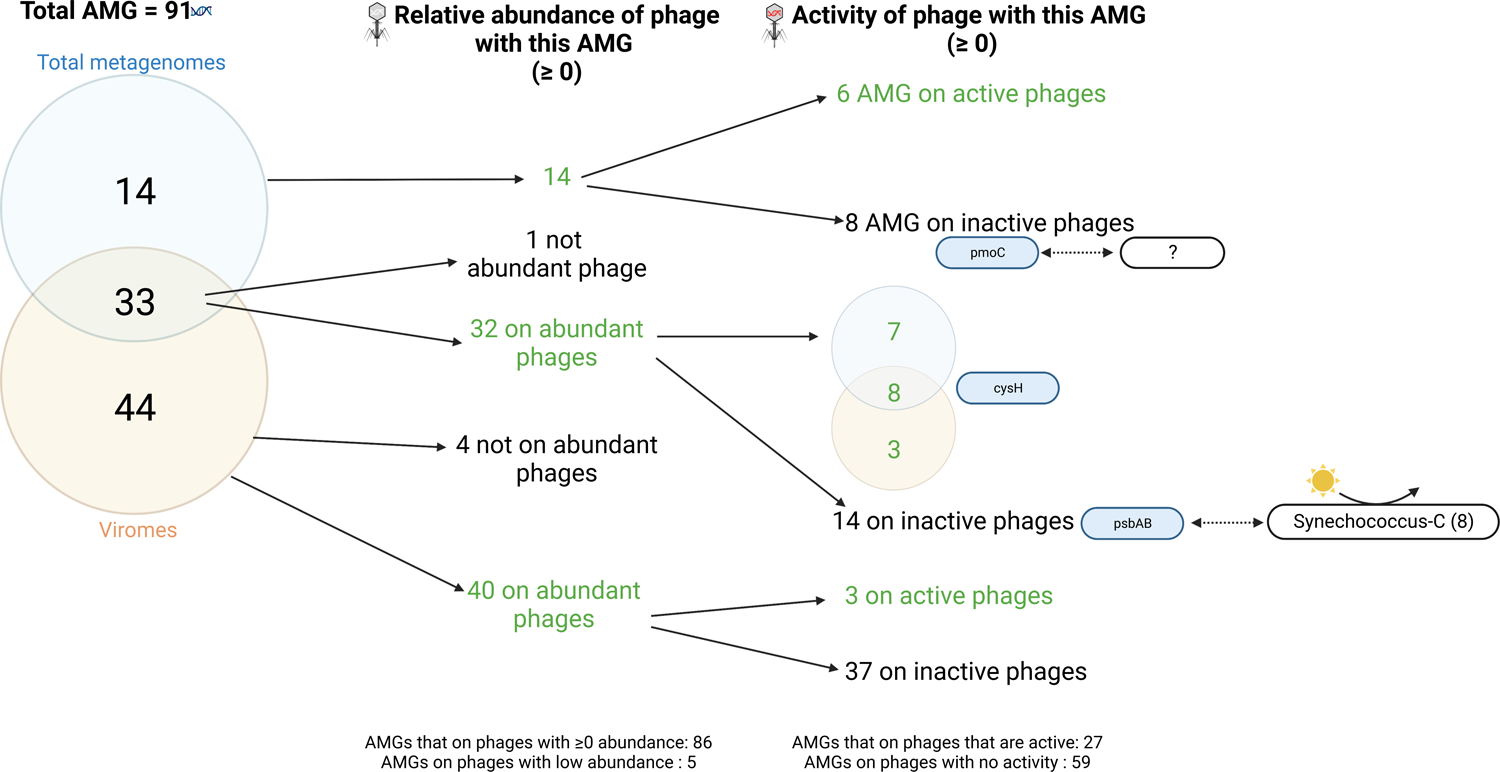
Phage and AMG expression, and examples of phage-host relationships. 91 auxiliary metabolic genes (AMGs) found on high-quality and complete phage genomes and their distribution between “expressed” and “not expressed”, and between total metagenomes and viromes. Of those, 59 AMGs were on phages that were not expressed, including ecologically relevant AMGs *pmoC* (aerobic methane oxidation), and *psbA* and *psbD* (photosynthesis). The *psbAD* AMGs were found on 2 phages whose hosts were 8 closely related species of Cyanobacteria, specifically *Synechococcus-C*. The *pmoC* AMG was on a phage that was not predicted to be linked to a specific prokaryote MAG in our dataset or to previously published bacterial and archaeal reference genomes.

### Phage-host interaction diversity and specificity

We predicted phage-host interactions by identifying the CRISPR sequences in host genomes, keeping in mind that approximately 22% of bacteria in a population maintain the signature of previous phage infection^29^. We identified 113 CRISPR cassette arrays in 68 MAGs of the 431 prokaryote MAGs, and 11 of those CRISPR arrays (in 9 MAGs) had a 100% match to phages (0 mismatches) with a Lake Mendota phage dataset (genomes of all qualities included), which assigned them to 22 phages. We inferred that an array that matched a phage corresponded to the susceptibility of that host to phage infection, at least at some point in the recent past. The inferred infected groups include Gammaproteobacteria, Bacteroidota, Cyanobacteria (*Dolichospermum flosaquae* and *Microcystis*), Verrucomicrobiota (Verrucomicrobiota and *Kirimatiellae*), Chloroflexota, and UBA2361. These phage were taxonomically assigned to *Caudoviricetes* (*Myoviridae*, *Siphoviridae*, and *Podoviridae*), although several could not be assigned. Two of the Chloroflexota phages encoded the *cysK* AMG, involved in cysteine metabolism. Most identified phages were obtained from virome samples as opposed to total metagenome samples. Each identified MAG appeared to be susceptible to a range of phages that are not closely related, except *Dolicospermum flosaquaea* which was infected by 5 taxonomically unknown phages sharing high ANI among themselves. In contrast, we found that phages displayed a high host-specificity (infected only 1 MAG) (**Supplementary Table 6**, **Supplementary Table 7**).

Additional ways to predict phage-host interactions rely on the host-virus similarity in sequences (e.g. tRNAs, prophages) and sequence composition (e.g. kmer) and viral marker genes indicative of hosts^29^. Machine learning approaches relying on features learned from reference databases of viruses with their known hosts was developed to assign phages to potential hosts, even without having host genomes. Therefore, to further identify phage-host relationships, we used iPHoP^30^, which integrated the above prediction methods to assign phages to putative hosts that may or may not have a MAG available. Among the 431 MAGs, 50 had a putative phage associated with them, 55 when including the additional manual CRISPR cassette method. When looking at the expression of those 50 MAGs across the 16 metatranscriptomes, 15 of the MAGs were never expressed. Among those expression categories, generally fewer MAGs were found to be associated with a phage across all categories. While a greater diversity of organisms was active in the hypolimnion, only those belonging to Gammaproteobacteria, Bacteroidota, and Actinobacteriota were directly impacted by phages.

### Phages impact biogeochemical processes driven by active microorganisms

To estimate the importance of these phage-host interactions on biogeochemical cycling, we predicted metabolism of the phage-impacted bacteria and mapped RNA-seq data onto these to determine activity (**Supplementary Table 8**). Among the 55 phage-associated MAGs, many hosts had key genes for aerobic methane oxidation, denitrification, nitrogen fixation, and sulfur cycling, many of which were also expressed (**Figure 6**). While we did recover a *Methylococcales* MAG that encoded all *pmoABC* genes for methane oxidation, its phage did not encode any homologous AMGs. Phage-associated *Burkholderiales*, Cyanobacteria, Fibrobacteria (inactive), and Verrucomicrobiota encoded genes for nitrogen fixation. *Burkholderiales*, especially *Rhodoferax* organisms had the full set of genes for complete denitrification. Additionally, several *Burkholderiales* organisms also possessed genes for sulfur oxidation and expressed them.

**Figure 6.**
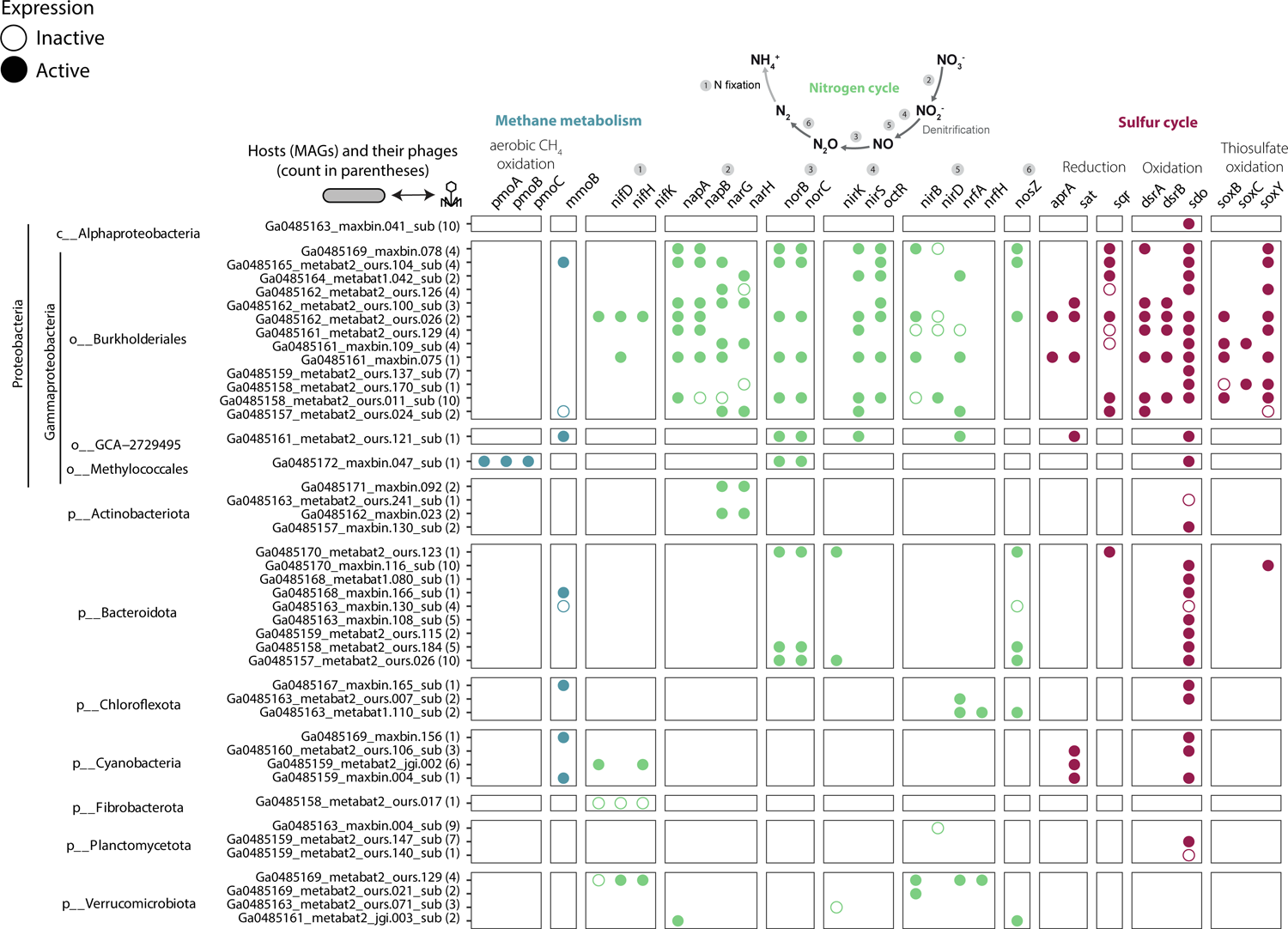
Methane, nitrogen, and sulfur metabolism and gene expression of 55 MAGs with at least one identified phage. 44 out of 55 MAGs had N, S or CH_4_ genes to be plotted. Phage susceptibility ranged from 1 to 10 phages per MAG. 1 MAG (*Methylococcales*) had all *pmoABC* genes in the aerobic methane oxidation pathway (the associated phage did not encode a pmo auxiliary metabolic gene. *Burkholderiales* (Gammaproteobacteria), Fibrobacteria, and Verrucomicrobiota MAGs had genes in nitrogen fixation. Phage-associated Gammaproteobacteria were involved in multiple steps of the denitrification and sulfur transformation pathways. No phage-associated host had genes in nitrification (aerobic ammonia oxidation nor nitrite oxidation). Note that some dsrD genes were found on some MAGs, but the gene was not active. Data from **Supplementary Table 8** was used for plotting.

### Phage-host abundance and expression dynamics

Any given phage was found between 1 and 16 samples and expressed (active) between 0 and 9 samples. Cyanophages were among the most present and active phages in the dataset. (**Supplementary Figure 5**) and included hosts from *Microcystis* (genera *Microcystis* and *Dolichospermum*) and *Nostocera* (10 different genera). Among all phages that were active, only 6 were present *and* active *and* had a binned MAG as a host (2 Cyanobacteria, 3 Bacteroidota, and 1 Gammaproteobacteria). The 3 Bacteroidota (Ga0485170_maxbin.116_sub, Ga0485159_maxbin.015, Ga0485168_metabat1.080_sub) were not among top-ranked abundant, active nor overexpressed MAGs. The phage-host abundance and expression patterns of the 2 Cyanobacteria MAGs and their cyanophages, and of the *Thiobacillacea* (Gammaproteobacteria) and its phage are shown in **Figure 7**. The first cyanobacteria MAG, the *Dolichospermum*, was abundant and active pre- and post-fall mixing, and phage-host dynamics were coordinated.

**Figure 7.**
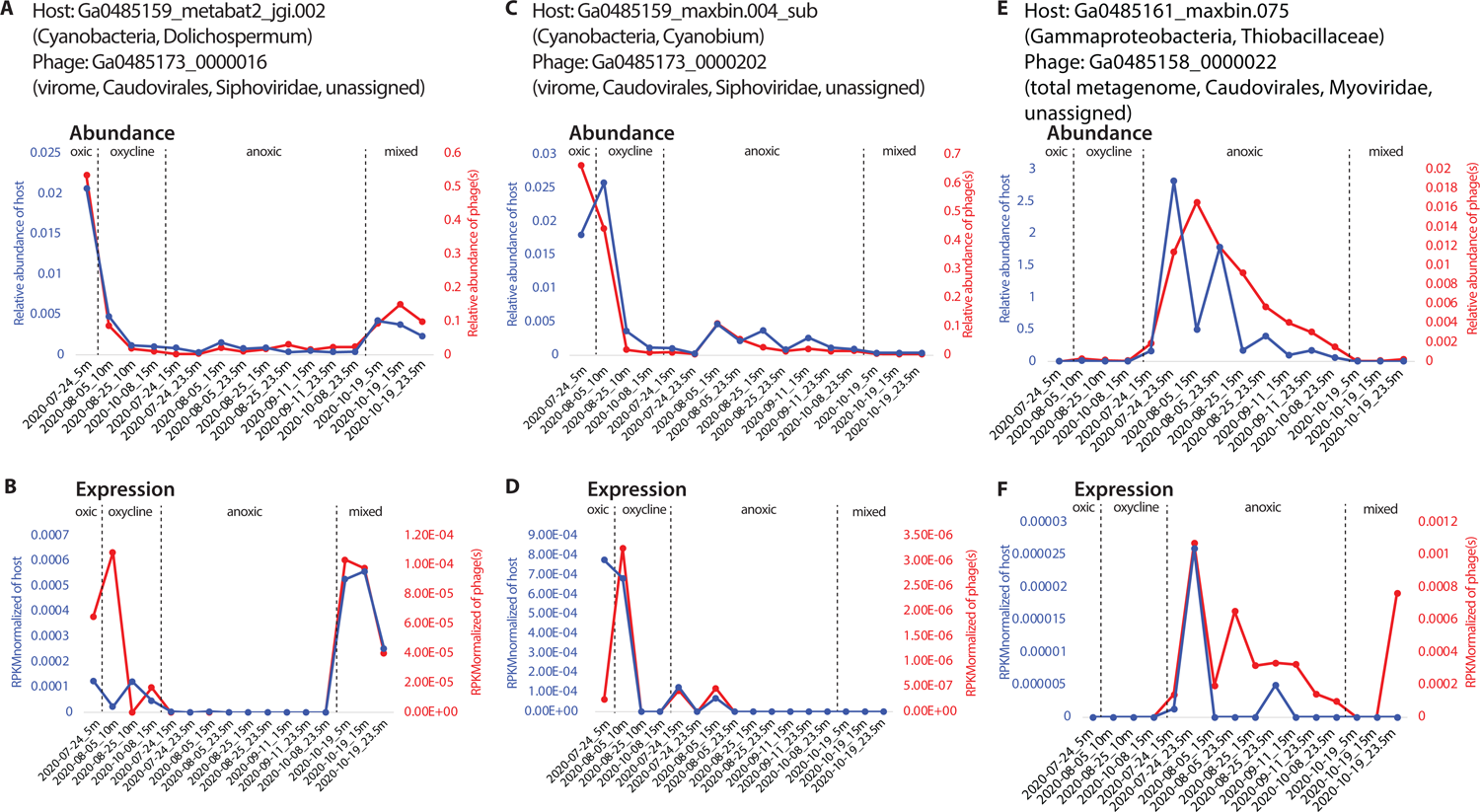
Phage-host abundance and activity patterns of Cyanobacteria and *Thiobacillaceae* hosts and their phages. **A and B.** The *Dolichospermum* phage-host pair’s abundance (A) and activity (B) show a higher abundance during stratification in the oxic epilimnion, and post mixing. In comparison, the phage was highly active in the oxycline and post mixing. This MAG is not among the most abundant ones but is in the top 10 ranked most active in 4 out of 16 samples, which correspond to the oxic samples (July 25 5m, and the 3 samples from Oct 19). **C. and D**. The *Cyanobium* phage-host pair’s abundance (C) and activity (D) show that these organisms are abundant in the upper water column and throughout the water column but were only active in the oxycline and in the upper anoxic zone (15m). This MAG is in the 7^th^ most active taxa in the July 5m (oxic) sample. **E and F.** This *Thiobacillaceae* ranked in the top 10 most abundant taxa of 5 anoxic samples and in the top 10 most active MAG in all the 9 anoxic samples and of the oxic-anoxic 23.5m sample (**Figure 2B** and **Supplementary Table 7**).

## Discussion

Here, we study how spatiotemporal impacts of stratification and anoxia influence microbial and viral communities and activity in a mid-size eutrophic lake, that is representative of temperate lakes worldwide. We investigated how biogeochemical cycles in seasonally anoxic freshwater environments were correlated with viruses and prokaryotes involved in those functions. Our analyses revealed differences in the abundance and expression levels of the prokaryotic community over weekly scales, mainly differentiated by stratification and oxygen levels, but this was not observed in the viral community. Additionally, our comparison of viromes and metagenomes revealed that viromes resulted in more high-quality and complete phage genomes and a higher number of previously undiscovered “unknown” phage taxa. Finally, we identified 9 MAGs with putative phages just based on CRISPR arrays, and a total of 55 MAGs with phages based on the integration of multiple phage-host prediction methods. These 55 MAGs represented 7 phyla. These bacterial hosts were inferred to be involved in methane oxidation, denitrification, nitrogen fixation, sulfur oxidation, and sulfate reduction, and many of these genes were indeed expressed. Next, we contextualize what is known about microbial communities of stratified anoxic lakes, freshwater phages, and the implications of this work beyond freshwater phage ecology.

### Increased microbial diversity over time and space is not directly correlated to activity

Prokaryotic diversity increased with time and water column depth. This trend might reflect the emergence of diverse and heterogenous microniches corresponding to the dynamics of carbon/nutrient availabilities and redox gradients with the succession of preferred terminal electron acceptors since the stratification. As the fall mix started, physicochemical “homogenization” of lake water occurred. For example, the entire water column became oxygenated, and as oxygen was the preferred electron acceptor thermodynamically, microbial processes with alternative electron acceptors were suppressed. As a result, such homogenization could “simplify” the microbial community, indicated by the decrease in Shannon diversity index post-mixing.

Environmental stratification and redox gradients result in strong differences in microbial community structures between the oxic epilimnion and the anoxic hypolimnion, and fall mixing can “reset” the water column’s microbial community^15^. Lake Mendota did not have many archaeal groups, unlike in anoxic freshwater lakes with deep hypolimnions^31, 32^. As for the epilimnion, in Lake Mendota, a historical time-series of 16S rRNA gene sequences revealed that the upper 12m of the water column, mostly oxygenated but sometimes spanning the oxycline, mostly contains Cyanobacteria, Actinobacteriota, Proteobacteria, and Bacteroidota^33^. In contrast, reconstructed genomes of Bacteroidota and the PVC superphylum (Planctomycetota, Verrucomicrobiota, Chlamydiota)^6^ were recoverd from the hypolimnion. 16S rRNA gene-amplicon-based studies of the microbial community across depths (2016-2017) revealed that taxonomic groups were strongly influenced by overall redox status, rather than the onset of anoxia^17^. In concordance with this, the changes in the oxycline depths across the season were related to the microbial community, and even in the hypolimnion, differences were observed between the 15m and 23.5m layers.

We noted that the abundance of taxa (based on metagenomic sequencing) did not necessarily correspond to the activity of taxa. For example, while the metagenomic data supported abundant groups commonly found in Lake Mendota and the freshwater lakes, our results highlight the importance of *Methylococcales*, Desulfobacteriota, and *Burkholderiales* as most active in the lake, especially in the oxycline and anoxic hypolimnion, which constitute a major ecological space during summer stratification.

### Viral impacts on biogeochemistry

Here, we specifically related phages to hosts (binned MAGs) from the same environment, in addition to comparing with viral host databases to identify putative hosts (not represented by our MAGs). The CRISPR-spacer similarity between hosts and viruses was widely used to link these pairs across environments, namely because these are natural bacterial and archaeal immune machinery against viral infections^34^. Consistent with the literature, phages displayed high host specificity, with one phage infecting one group of bacteria^35^. In other systems, ∼2000 phages were associated with 90 MAGs across 25 phyla, in a community with a total of 29 phyla^36^. Here, our initial overall community is less diverse (27 phyla). We found phages infecting Cyanobacteria, Gammaproteobacteria, Alphaproteobacteriota, Actinobacteriota, Fibrobacteria, Verrucomicrobiota, Bacteroidoita, Chloroflexota, and Planctomycetota – all are common freshwater phyla. Cyanophages have been reported in freshwater lakes, and other commonly found freshwater taxa such as Actinobacteria, Alphaproteobacteria, Planctomycetes, and Verrucomicrobiota have been previously reported to be infected by phages^37, 38^. We also found phages infecting Chloroflexota and Gammaproteobacteria (orders *Burkholderiales*, *Methylococcacea*, and GCA_2729495). In lakes, Chloroflexota is often associated with hypolimnion^39^ and can make up to abundance up to ∼25% in lakes^40^. Gammaproteobacteria phages have also been found in the Baltic Sea^41^, and *Methylococcaccea* phages have been found in Lake Baikal^13^.

Another case is when we found phages, but no host could be found. Methodological reasons may explain this, as unbinned scaffolds (up to 50% of the metagenome sometimes) could contribute to incomplete host genome(s). In such cases, leveraging public databases enabled us to identify potential hosts: for example, we found photosynthesis AMGs on phages infecting *Synechococcus* species when expanding our search to the reference database. Additionally, while we reconstructed multiple Cyanobacteria genomes, we did not resolve a *Synechococcus* genome, but did recover closely-related *Cyanobium* species (9 MAGs). *Synechococcus* spp. are known to be found in freshwater lakes though, for example in oligotrophic reservoirs^44^, along with their cyanophages^45^. In these cases, we can only infer the potential effect of the phage on the biogeochemistry of the lake – *Synechococcus* is among the most well-studied oxygenic photosynthetic bacteria, but this inference about the ecological role based on the host might be less obvious for other taxa.

Additionally, *Methylococcales* (Ga0485172_maxbin.047_sub) encoding *pmoABC* genes for aerobic methane oxidation, were among the most expressed MAGs in our dataset. Denitrification in the anoxic hypolimnion results in a decrease of NO_3_^-^ and an accumulation of intermediates. As for sulfate, about 45% of the sulfide in the sediment is derived from sulfate reduction in Lake Mendota^21^, and prior studies have shown that sulfate reduction could occur in the water column based on MAGs and biogeochemical profiles^6^. Our results support these findings, as many sulfate-reducing bacteria were active based on metatranscriptomic data, for example, the *Desulfobacula* genus within the *Desulfobacteraceae* family which are known to be sulfate-reducers.

Capturing active MAGs and active phages also depends on the timing of sampling. It has been shown that environmental factors such as salinity, temperature, UV-light^42^, and the host growth rate^43^ can influence phage communities and lifestyles, including the switch from lysogenic to lytic. In freshwater lakes, these environmental parameters change over time and space, sometimes following daily, weekly, seasonal, annual, or even decadal patterns.

### Collecting viromes in conjunction with total metagenomes allows for recovering more diversity of phages

Compared to cell-associated (total) metagenomes, a higher number of phages were recovered from viromes, supporting similar observation in soils^14^. In freshwater lakes, sequencing depth^37^ and different size fractions (based on the filter pore size)^46^ have been shown to influence which viruses are detected, as such it was beneficial to pursue paired metagenome-viromes in this study to capture the entire microbial community. Here, we found more lytic phage genomes than lysogenic. This goes against our expectation that metagenomes would recover more lysogenic phages, and this might have been caused by free lytic viral particles that would otherwise pass through the filter and could accumulate on the filter as it was clogged with biomass accumulated during the filtration process.

The impact of collecting and analyzing viromes has been demonstrated in oceans^47, 48^ and laboratory procedures have been optimized^49^. In freshwaters, for example, viruses were extracted from total cellular metagenomes in multiple lakes ^9, 13, 50, 51^. Viromes have been generated for freshwater lakes^52^, with varying sequencing methods, and paired with either metagenomes or transcriptomes using the technologies that were current at the time of those studies. Many physical separation methods of viruses have also been used in the past, including tangential flow filtration^53^, FeCl_3_ precipitation^54^ and PEG^55, 56^. Our combined total cellular metagenomes, viromes, and metatranscriptomes provide further evidence of not only the community structure and composition but also the activity of those populations in freshwater lakes.

Knowing that phages are generally highly specific to their hosts and that their hosts are influenced by environmental factors (e.g stratification and mixing), we also hypothesized that the viral community in the hypolimnion would be distinct from that of the epilimnion. This was not observed, instead, sampling methodology was a greater contributor to community differences. Recently, phages have been suggested to have a broader range of potential hosts, even domains^57^, potentially explaining why we observed less-specific phage-host impact than anticipated. Another implication of this work was the high proportion of “viral dark matter”^58^ in the virome fraction. The identifiable viral fraction of Lake Mendota was dominated by the *Caudoviricetes* order (*Podoviridae*, *Myoviridae*, and *Siphoviridae*), as are many other freshwater lakes^7, 53, 54, 59–62^. The high-quality, complete viral genomes reported here will expand the repertoire of freshwater phage genomes^59^, which we anticipate becoming a genomic resource for the microbial ecology research community, enabling cross-site comparisons in the future.

### Careful interpretation of phage AMG and their actual activity is needed

While several phage genomes were obtained, and many AMGs were found with functions ranging from core cellular metabolism to biogeochemically relevant pathways, improved ways to interpret the data remain an urgent need. Here, we observed up to 4.5% of all recovered complete and high-quality phage genomes were expressed at any given point in time. Even fewer were captured as expressing AMGs using RNA-seq. For example, while many AMGs were encoded by phages from both total metagenome and virome fractions (e.g. *psbA*, *psbD* in photosynthesis), many AMGs were sometimes only found in the total metagenomes (e.g. *pmoC* involved in methane and ammonia oxidation) or only in viromes (e.g. *soxB*). Further, capturing phage activities from metatranscriptomes could be a matter of luck, to sample and sequence the exact “time snapshot” when the lytic phase occurs. Few AMGs were expressed at any given sampling time, with most phages encoding core metabolism AMGs being the most expressed. These findings imply that depending on the research question and the biogeochemical cycle of interest, ecological interpretation from total metagenomes can over- or under-estimate true activity. This must be kept in mind when analyzing viral scaffolds from total metagenomes and inferring activity. For example, the lack of AMGs in a cycle of interest, and the lack of viruses with an identifiable host may be a result of a methodological bias (false negative), rather than true absence, and similarly, the presence of an AMG of interest may be in an inactive phage and inactive host.

Unlike for prokaryotic community identification, viral identification remains strongly impacted by laboratory methods. Methods prior to sequencing DNA impacted the viral community composition observed, functional genes found, and therefore reference genomes onto which transcriptomes were mapped onto to determine activity. These findings are applicable beyond freshwater phage ecologists, as anyone working in viromics data and using bioinformatics tools to bring biological insight should be careful about introduced biases or acknowledge interpretation caveats. Second, we found that both prokaryotic and phage activity were not correlated with abundance, which implies that abundance-only or genomic potential methods can lead to overestimation or underestimation of activity. Finally, as environmental phage detection becomes applied to water quality, for example in the context of public health^63^, and bioinformatics tools are developed to make sense of complex genomic data, careful attention should be given to the way that available datasets (mostly bulk metagenomes, not viral-fraction specific, or reference-based viral inference) impact detection and interpretation.

## Methods

### Field sampling

Lake Mendota, Wisconsin, USA was sampled at the “Deep Hole” location (Latitude 43°05’54”, Longitude 89°24’28”), during the open-water period from July 7, 2020, to October 19, 2020. The samples were collected approximately weekly at 5 depths (5, 10, 15, 20, and 23.5m) and spanned the shift from a stratified water column with oxic, oxycline, and anoxic layers to a mixed oxygenated water column on October 18, 2020 (**Figure 1**).

Water was filtered back in the lab (less than 1-5 hours after sampling) through a serial vacuum pump through a 0.22μm filter to capture cells for total metagenome or metatranscriptome sequencing. The filters were stored at −80°C until DNA or RNA could be extracted. The filtrate from 5, 15, and 23.5m was further amended with FeCl_3_, a flocculation technique that concentrates virions for the viromes. The FeCl_3_ protocol was adapted from^64, 65^. In short, FeCl_3_ stock was made, and 1000uL was added to ∼500mL of lake water filtrate. Bottles were manually mixed for 5 minutes and left at room temperature to incubate for 1 hour. Then FeCl_3_-amended lake was filtered through a 0.8μM filter to collect the viral biomass.

### DNA extraction

We sequenced 30 samples (16 total metagenomes and 14 viromes) through the US Department of Energy Joint Genome Institute (JGI). We chose those 30 samples out of the 200+ filters collected based on their position in the water column (i.e. whether they were in oxic, oxycline, or anoxic portions of the lake) (**Supplementary Table 1**), as well as to capture the pre- and post-fall mixing microbial community. The DNeasy PowerSoil kit (Qiagen) was used to extract DNA from both the >0.22 μm and <0.22 μm fractions. The DNA was quantified using the QuBit DNA.

### RNA extraction

RNA extraction was performed using a phenol-chloroform protocol on 16 filters that corresponded to the 16 total metagenomes^6, 66^. In summary, frozen filters were put into a bead-beating tube (e.g. PowerSoil (Qiagen)), and 1mL of TRIzol (Invitrogen) was added to each filter. After the filters were bead beaten for 1 minute at the medium speed, an internal standard (i.e. an *in vitro* transcribed pFN18A of a known concentration) was added^6, 66^ and inverted to mix. The samples were centrifuged for 1 min. The supernatant was transferred and 300uL of chloroform was added, mixed by inversion for 1 min, and incubated at room temperature for 3 minutes. After 15 min of centrifugation at 4°C (13200xg), the supernatant was transferred to a new tube, and cold 70% ethanol was added and mixed well. Next, the RNeasy extraction kit (Qiagen) was used to recover the extracted RNA. DNase treatment was performed using the TURBO DNA-free™ Kit (Invitrogen), which contains the DNase inactivation reagent. RNA was quantified using the QuBit RNA kit (Invitrogen).

### Biogeochemical field data collection and laboratory analysis

Due to the COVID-19 pandemic, sampling and collecting of field data were limited. While we initially planned on collecting weekly sulfate, sulfide, nitrate, and nitrite data for 2020, we were unable to collect all biogeochemical data. Therefore, in 2021, we collected sulfate and sulfide. To collect sulfide in the field, 9 mL of lake water was fixed with 1 mL of 10% Zinc Acetate, mixed, then stored in a cooler with dry ice for sulfide measurements. Zinc acetate addition prevents the oxidation of sulfide with oxygen. In the lab, we used the Cline assay to measure sulfide.

For sulfate data collection, we collected ∼10 mL of unfiltered lake water and froze it with dry ice in the field. Samples were brought back to the lab and immediately stored at −20°C until processing. Sulfate data was analyzed using ion-chromatography (IC) (Dionex 4mm ICS-2100 anion analyzer). Five standards were made with concentrations ranging from 0.8ppm −80ppm of sulfate (MgSO_4_ was used as the source of sulfate). 7 ml of each standard and sample were used form measurements. The loop size was 25μL, and the ion-chromatography columns used were AGII-HC and ASII-HC, the mobile phase was 30mM NaOH, and the suppressor was ASRS-4mm. One standard was checked every 10 samples to ensure proper measurements throughout. Results were analyzed in the software Chromeleon.

### DNA sequencing

16 total metagenomes and 14 viromes were sequenced at the Joint Genome Institute (JGI) (**Supplementary Table 1**). The JGI Gold Project ID is Gs0151897, under the name “Freshwater bacterial and viral communities from Lake Mendota, Wisconsin, United States”. Details of sequencing for each sample can be found on the JGI portal, using the “Report” tab.

For the 16 total metagenomes, plate-based DNA library preparation for Illumina sequencing was performed on the PerkinElmer Sciclone NGS robotic liquid handling system using the Kapa Biosystems library preparation kit. ∼2 ng of sample DNA was sheared to 506 bp using a Covaris LE220 focused-ultrasonicator. The sheared DNA fragments were size selected by double-SPRI and then the selected fragments were end-repaired, A-tailed, and ligated with Illumina compatible sequencing adaptors from IDT containing a unique molecular index barcode for each sample library. The prepared libraries were quantified using KAPA Biosystems’ next-generation sequencing library qPCR kit and run on a Roche LightCycler 480 real-time PCR instrument. The flowcell was sequenced on the Illumina NovaSeq sequencer using NovaSeq XP V1.5 reagent kits, S4 flowcell, following a 2×151 indexed run recipe.

For the 14 viromes, ∼0.02 ng of DNA was denatured and sheared to ∼300 bp using the Covaris LE220, and >300 bp was size selected using SPRI beads (Beckman Coulter). The fragments were denatured again and a library was created using Accel-NGS 1S Plus DNA Library Kit (Swift Biosciences) to ligate Illumina-compatible adapters (IDT, Inc). qPCR was used to determine the concentration of the libraries which were sequenced on the Illumina platform. The prepared libraries were quantified using KAPA Biosystems’ next-generation sequencing library qPCR kit and run on a Roche LightCycler 480 real-time PCR instrument. The flowcell was sequenced on the Illumina NovaSeq sequencer using NovaSeq XP V1.5 reagent kits, S4 flowcell, following a 2×151 indexed run recipe.

### RNA sequencing and processing

RNA sequencing was performed at SeqCenter (PA). Illumina Stranded RNA library preparation with RiboZero Plus rRNA depletion to offer RNA Sequencing. Metatranscriptomes were quality-trimmed using fastp with a quality score cutoff of 20. The following non-coding rRNA reads were removed using sortmerna (v4.2.0)^67^: 5.8S and 5S based on RFAM database^68^; archaeal and bacterial 16S and 23S based on SILVA database (v.119); and eukaryotic 18S and 28S based on SILVA database (v.119)^69^. Reads mapping to the internal standard were then identified using sortmerna and quantified. Reads not associated with the internal standard or rRNA were used for further analysis.

### Metagenome assembly, binning, and MAG quality filtering

The quality of the assembled scaffolds, assembled with SPADES v.3.13.0^70^ with the following options: -m 2000 -o spades3 --only-assembler -k 33,55,77,99,127 –meta) was assessed by 1) quantifying the percentage of original reads mapping back to the scaffolds, with a value of >80% meaning a good assembly quality 2) the distribution of the lengths of the scaffolds in the assembly and 3) the N50 value of the assembly. This information is found in the PDF report from JGI available on IMG/M.

A complete workflow for the metagenomic binning is shown in **Supplementary Figure 6**. Binning was performed using Metawrap v1.3.2^71^, a binning wrapper that relies on Metabat1 v0.32.5^72^, Metbat2 v2.12.1^73^, Maxbin2^74^, using default settings, but using differential coverage generated by mapping reads from each of the 16 metagenomes to each of the 16 assemblies. Cross-mapping here means that all 16 interleaved fastq files were used to bin each 16 total metagenomes. We compared 3 different binning methods: Binning using reads from itself, binning using reads from samples from the same ecosystem (oxic, anoxic-oxic, and anoxic depths) “within layer”, and cross-mapping across all samples. We chose cross-mapping for all samples because compared to binning using the respective assembly only *or* binning with reads from the same environmental layer, cross-mapping resulted in the highest number of bins *and* the highest average bin completeness (**Supplementary Figure 7**). Further, to improve bin quality and completeness, we next performed bin refinement.

Bin refinement was performed using DAS Tool v.1.1.3^75^. Bin refinement involves clustering bins that may be from the same population and uses algorithms to build “better”, more complete bins^75^. The bins provided for each sample came from metabat1, metabat2, maxbin (ran through Metawrap) using differential coverage, and a set of independently generated bins from JGI, which uses metabat2 but without the differential coverage (that is 1 fastq file was used with each 1 assembly during the metabat2 command). These differences in metabat2 versions and settings used are why we provided all “four” binning results as input for bin refinement. After bin refinement, on average, 15 to 20% of bins were kept (1,445 MAGs).

We then performed a taxonomic assignment using GTDB-tk v.1.7.0^76^. This was done early, as opposed to after getting the final bin set, because the dereplication step relies on genome quality estimate calculation by CheckM v1.1.3^77^. CheckM can be run without default marker genes, or with a CPR (i.e. Patescibacteria) marker gene set, which results in more accurate estimates for CPR genomes. By running the taxonomic assignment first, we will divide the refined bin set into CPR genomes and non-CPR genomes, and then run CheckM with the appropriate marker gene set accordingly. In total, 43 CPR bins, and 1,402 non-CPR bins (17 of which were Archaea).

To reduce redundancy in the MAGs across the 16 metagenome samples, dereplication was done using dRep v.3.2.2^78^. Unlike bin refinement which improves upon existing bins and may create “new” bins from the inputs, dereplication clusters bins and selects one representative bin from each cluster. As we move through the processing of the metagenomic data, it is always possible to go back, and even though only 1 representative genome is selected after dereplication, it is possible to know which other very closely related genomes existed, which can be useful for projects outside the scope of this paper. The completeness ≥50% and contamination <=10% thresholds were used as they generally represent MIMAG standards^79^. After dereplication, we obtained 18 CPR MAGs and 413 non-CPR MAGs (2 were Archaea) for a total of 431 MAGs. These 431 MAGs consisted of the “final bin set”.

### MAG naming convention

MAGs from the final bin set are described in **Supplementary Table 2**. The naming pattern is as follows: “GaXXXXXXX” represents the GaID of the metagenome or viromes, searchable on the JGI IMG/M website, from which the bin was originally generated. Following are “metabat1”, “metabat2_ours”, or “maxbin2” representing the binning tool used, all with differential coverage. If the name has “_jgi”, the MAG was obtained using metabat2 without differential coverage, as originating from the standard JGI sample processing pipeline. Following is a number with 3 digits, representing an arbitrary bin number within each binning run. If the bin name has “_sub” in it, it means the MAG results from bin refinement (and improved upon the original binning”. Because the final bin set represents closely related MAGs (dereplicated set), the GaID of the MAG does not mean the MAG is from that sample only, instead that the best representative MAG originated from it.

### Assessing genome coverage and transcript abundance per MAG

We used CoverM^80^ to obtain genome-level relative abundance and expression for all 431 MAGs and phages. We used CoverM with the option -m rpkm to assess the expression of the 431 MAGs across all the samples. Because we used internal standards, we were able to get a read count of the internal standard for each of the 16 samples. We divided the rpkm by the respective internal standard read count to obtain the “rpkm normalized” value. To calculate the genome-level coverage, we used Bowtie2 v.2.2.5^81^ with default settings to build an index of the 431 MAGs, sorted it with samtools sort v.1.14^82^, and then used inStrain v.1.5.7^83^ quick_profile to obtain per-genome depth and breadth of the coverage.

### Assessing the metabolic potential of the MAGs

We used METABOLIC v.4.0 ^84^ on all 431 MAGs to get an overview of genes involved in carbon, nitrogen, sulfur cycling, and in general core metabolism (KEGG^85^, Pfam^86^, CAZyme^87^, MEROPS^88^).

### Phylogenetic tree

A concatenated gene phylogeny of 16 ribosomal proteins from the 431 MAGs was created to gain an overview of the taxonomic diversity in Lake Mendota, and visualize the distribution of the organisms over time, and any metabolic genes of interest. The function “create-gene-phylogeny” in MetabolisHMM v. 2.21^89^ was used to create a bacterial tree of 429 MAGs (the 2 archaeal genomes were not included as different marker genes are used when building an archaeal tree and a bacterial tree, and because a 2-genome tree is not very informative). The *fasta* alignment file from metabolisHMM was then used as an input to generate a maximum-likelihood phylogenetic tree using RAxML Blackbox HPC v.8.2.11^90^, through the CIPRES server v.3.3. iTOL (interactive Tree of Life)’s interface was used to visualize and annotate the tree.

### Diversity Index Calculations

The Shannon Diversity Index and the beta-diversity index were calculated in R using the package “vegan” v.2.5.7 with an abundance matrix grouped at the class level.

#### Viral identification and AMG annotation

VIBRANT^91^ was used on the assembled scaffolds of the 16 metagenomes and the 14 viromes to identify viral genomes, and to identify whether the phage was likely lytic (non-integated) or lysogenic. The -virome option was used on the viromes. In total, about 500,000 phages were found, representing about 4 million protein-coding sequences. Phage genome quality was assessed with CheckV^92^.

#### Phage-host interactions

The method of host homology was initially used to assess sequence similarity between the 431 MAGs and all phages that we recovered from Lake Mendota using DIAMOND blastp^93^. We first identified CRISPR sequences in the 431 MAGs to narrow down interactions, as CRISPR sequences in bacterial hosts represent the phages that may infect them. CRISPR sequences were identified with piler-cr^94^, to identify repeats and spacers. To link MAGs containing spacer sequences to phages that may infect, we used BLAST homology of identified CRISPR spacers vs. all the Lake Mendota phages, allowing for at most 1 mismatch. To further attempt to assign phage-host interactions, especially for phages whose hosts are not represented by MAGs, we selected the 2,246 complete and high-quality phages in our study and ran iPHoP^30^ to assign putative hosts using the regular iPHoP database, with confidence scores greater or equal to 90. We also ran the search on a custom database by adding the 431 MAGs from our study to the regular iPHoP (*add_custom_db* option).

#### Taxonomic assignment of phages

We used Diamond blastp against the NCBI taxonomic database to assign taxonomy to the phages.

### Data Availability

The data is available at NCBI BioProject PRJNA758276, which contains all 431 prokaryotic metagenome-assembled genomes and the processed 16 RNA-seq samples. The phage genomes can be accessed at https://doi.org/10.6084/m9.figshare.22213846. The original and assembled metagenomes were sequenced through the JGI Gold Gs0151897, and the sequenced data and assembled scaffolds can be downloaded on JGI IMG/M. The R code is available at: https://github.com/patriciatran/LakeMendota-PairedOmics

## Acknowledgments

We are grateful to the Center for Limnology and the Long-Term Ecological Research – North Temperate Lakes group for supporting field sampling. PQT received funding from the Natural Science and Engineering Research Council of Canada (NSERC) doctoral fellowship, the Anna Grant Birge Memorial Award from the Center for Limnology, and the Baldwin Distinguished Graduate Fellowship from the Department of Bacteriology at the University of Wisconsin-Madison. DNA sequencing was conducted at the DOE Joint Genome Institute, a DOE Office of Science User Facility, via a Community Science Program New Investigator award to KA and PQT (award number 506328). KA acknowledges support from the USDA National Institute of Food and Agriculture (NIFA) under grant: Hatch project 1025641, and from the National Science Foundation grant DBI-2047598.

## Competing interests

The authors declare no competing interests.

## Supplementary Figures and Legends

**Supplementary Figure 1.** Sulfate and sulfide profiles (μM) in Lake Mendota in 2019 and 2021.

**Supplementary Figure 2.** Phylogenetic tree of the 429 Bacterial MAGs using 16 ribosomal proteins. The 2 Archaea are not shown.

**Supplementary Figure 3.** Bar plot showing the abundance and expression of all MAGs in Lake Mendota in this study.

**Supplementary Figure 4.** Bar plot showing the taxonomic composition of the phages in Lake Mendota in this study.

**Supplementary Figure 5.** Number of phages and in how many samples are they expressed or present in.

**Supplementary Figure 6.** Metagenomic processing workflow.

**Supplementary Figure 7.** Comparison of binning using individual sample reads only, reads from samples with similar environmental parameters (e.g. oxygen levels), or differential coverage using all samples. Metabat2 with cross-mapping resulted in an overall higher number of MAGs (A), but maxbin2 with cross-mapping resulted in the highest overall quality of MAGs (B). For this reason, cross-mapping was chosen as the method to obtain more, and high-quality MAGs, before refinement and dereplication.

## Supplementary Tables and Legends

**Table S1.** Sample description

**Table S2.** Genome characteristics for the 431 MAGs, including CheckM and GTDB-tk results.

**Table S3.** Genome-wide average-nucleotide identity (% ANI) comparison with previously published MAGs from Lake Mendota.

**Table S4.** Phage genome quality characteristics and sizes, in the total metagenomes and viromes.

**Table S5.** Abundance and expression of the high-quality and complete phages, with labels for whether they encoded auxiliary metabolic genes.

**Table S6.** Host susceptibility of MAGS

**Table S7.** Phage specificity.

**Table S8.** Presence and absence of genes in the 431 MAGs, along with whether these genes were expressed (i.e. active at any point in the dataset).

## Notes

### Competing Interest Statement

The authors have declared no competing interest.

## References

1. Snortheim, C. A. et al. Meteorological drivers of hypolimnetic anoxia in a eutrophic, north temperate lake. Ecological Modelling 343, 39–53 (2017).

2. Schwefel, R., Gaudard, A., Wüest, A. & Bouffard, D. Effects of climate change on deepwater oxygen and winter mixing in a deep lake (Lake Geneva): Comparing observational findings and modeling: CLIMATE CHANGE EFFECTS IN A DEEP LAKE. Water Resources Research 52, 8811–8826 (2016).

3. Jane, S. F. et al. Widespread deoxygenation of temperate lakes. Nature 594, 66–70 (2021).

4. Linz, A. M. et al. Freshwater carbon and nutrient cycles revealed through reconstructed population genomes. PeerJ 6, e6075 (2018).

5. Jones, D. S. et al. Molecular evidence for novel mercury methylating microorganisms in sulfate-impacted lakes. ISME J 13, 1659–1675 (2019).

6. Peterson, B. D. et al. Mercury Methylation Genes Identified across Diverse Anaerobic Microbial Guilds in a Eutrophic Sulfate-Enriched Lake. Environ. Sci. Technol. 54, 15840– 15851 (2020).

7. Sime-Ngando, T. et al. Short-term variations in abundances and potential activities of viruses, bacteria and nanoprotists in Lake Bourget. Ecol Res 23, 851–861 (2008).

8. Breitbart, M., Bonnain, C., Malki, K. & Sawaya, N. A. Phage puppet masters of the marine microbial realm. Nature Microbiology 3, 754–766 (2018).

9. Roux, S. et al. Ecogenomics of virophages and their giant virus hosts assessed through time series metagenomics. Nature Communications 8, (2017).

10. Djikeng, A., Kuzmickas, R., Anderson, N. G. & Spiro, D. J. Metagenomic analysis of RNA viruses in a fresh water lake. PLoS One 4, e7264 (2009).

11. Langlois, V., Girard, C., Vincent, W. F. & Culley, A. I. A Tale of Two Seasons: Distinct Seasonal Viral Communities in a Thermokarst Lake. Microorganisms 11, 428 (2023).

12. Potapov, S. A. et al. Metagenomic Analysis of Virioplankton from the Pelagic Zone of Lake Baikal. Viruses 11, (2019).

13. Coutinho, F. H. et al. New viral biogeochemical roles revealed through metagenomic analysis of Lake Baikal. Microbiome 8, 163 (2020).

14. Santos-Medellin, C. et al. Viromes outperform total metagenomes in revealing the spatiotemporal patterns of agricultural soil viral communities. ISME J 15, 1956–1970 (2021).

15. Shade, A., Jones, S. E. & McMahon, K. D. The influence of habitat heterogeneity on freshwater bacterial community composition and dynamics. Environmental Microbiology 10, 1057–1067 (2008).

16. Rohwer, R. R., et al. The aftermath of a trophic cascade: Increased anoxia following species invasion of a eutrophic lake. http://biorxiv.org/lookup/doi/10.1101/2023.01.27.525925 (2023) doi:10.1101/2023.01.27.525925.

17. Marick, R. A., Peterson, B. D. & McMahon, K. D. Stratification in Microbial Communities with Depth and Redox Status in a Eutrophic Lake Across Two Years. http://biorxiv.org/lookup/doi/10.1101/2021.10.15.464574 (2021) doi:10.1101/2021.10.15.464574.

18. Lathrop, R. C. Perspectives on the eutrophication of the Yahara lakes. Lake and Reservoir Management 23, 345–365 (2007).

19. Nriagu, J. O. SULFUR METABOLISM AND SEDIMENTARY ENVIRONMENT: LAKE MENDOTA, WISCONSIN. Limnology and Oceanography 13, 430–439 (1968).

20. Magee, M. R. & Wu, C. H. Response of water temperatures and stratification to changing climate in three lakes with different morphometry. Hydrology and Earth System Sciences 21, 6253–6274 (2017).

21. Brock, T. D. Long-term Change in Lake Mendota. in A Eutrophic Lake: Lake Mendota, Wisconsin (ed. Brock, T. D.) 203–215 (Springer, 1985). doi:10.1007/978-1-4419-8700-6_9.

22. Kodama, Y. & Watanabe, K. Sulfuricurvum kujiense gen. nov., sp. nov., a facultatively anaerobic, chemolithoautotrophic, sulfur-oxidizing bacterium isolated from an underground crude-oil storage cavity. Int J Syst Evol Microbiol 54, 2297–2300 (2004).

23. Muck, S. et al. Niche Differentiation of Aerobic and Anaerobic Ammonia Oxidizers in a High Latitude Deep Oxygen Minimum Zone. Frontiers in Microbiology 10, (2019).

24. Roux, S., Matthijnssens, J. & Dutilh, B. E. Metagenomics in Virology. in Encyclopedia of Virology 133–140 (Elsevier, 2021). doi:10.1016/B978-0-12-809633-8.20957-6.

25. Chen, L.-X. et al. Large freshwater phages with the potential to augment aerobic methane oxidation. Nat Microbiol 5, 1504–1515 (2020).

26. Edwards, R. A., McNair, K., Faust, K., Raes, J. & Dutilh, B. E. Computational approaches to predict bacteriophage–host relationships. FEMS Microbiol Rev 40, 258–272 (2016).

27. Roux, S. et al. iPHoP: an integrated machine-learning framework to maximize host prediction for metagenome-assembled virus genomes. 2022.07.28.501908 Preprint at https://doi.org/10.1101/2022.07.28.501908 (2022).

28. Ingvorsen, K. & Brock, T. ELECTRON FLOW VIA SULFATE REDUCTION AND METHANOGENESIS IN THE ANAEROBIC HYPOLIMNION OF LAKE MENDOTA. LIMNOLOGY AND OCEANOGRAPHY 27, 559–564 (1982).

29. Fallon, R., Harrits, S., Hanson, R. & Brock, T. THE ROLE OF METHANE IN INTERNAL CARBON CYCLING IN LAKE MENDOTA DURING SUMMER STRATIFICATION. LIMNOLOGY AND OCEANOGRAPHY 25, 357–360 (1980).

30. Diao, M., Huisman, J. & Muyzer, G. Spatio-temporal dynamics of sulfur bacteria during oxic--anoxic regime shifts in a seasonally stratified lake. FEMS Microbiology Ecology 94, (2018).

31. Dolhi, J. M., Ketchum, N. & Morgan-Kiss, R. M. Establishment of microbial eukaryotic enrichment cultures from a chemically stratified Antarctic lake and assessment of carbon fixation potential. Journal of visualized experiments 1–5 (2012) doi:10.3791/3992.

32. Kankaala, P., Huotari, J., Peltomaa, E., Saloranta, T. & Ojala, A. Methanotrophic activity in relation to methane efflux and total heterotrophic bacterial production in a stratified, humic, boreal lake. Limnology and Oceanography 51, 1195–1204 (2006).

33. Robert Hamersley, M., et al. Water column anammox and denitrification in a temperate permanently stratified lake (Lake Rassnitzer, Germany). Systematic and Applied Microbiology 32, 571–582 (2009).

34. Juottonen, H. et al. Archaea in boreal Swedish lakes are diverse, dominated by Woesearchaeota and follow deterministic community assembly. Environmental Microbiology 22, 3158–3171 (2020).

35. Callieri, C. et al. The mesopelagic anoxic Black Sea as an unexpected habitat for Synechococcus challenges our understanding of global “deep red fluorescence”. The ISME Journal 13, 1676–1687 (2019).

36. Tran, P. Q. et al. Depth-discrete metagenomics reveals the roles of microbes in biogeochemical cycling in the tropical freshwater Lake Tanganyika. ISME J 15, 1971–1986 (2021).

37. Newton, R. J. & McMahon, K. D. Seasonal differences in bacterial community composition following nutrient additions in a eutrophic lake. Environmental Microbiology 13, 887–899 (2011).

38. Species invasions shift microbial phenology in a two-decade freshwater time series | bioRxiv. https://www.biorxiv.org/content/10.1101/2022.08.04.502871v1.abstract.

39. Zhang, M., Zhang, T., Yu, M., Chen, Y.-L. & Jin, M. The Life Cycle Transitions of Temperate Phages: Regulating Factors and Potential Ecological Implications. Viruses 14, 1904 (2022).

40. Williamson, S. J. & Paul, J. H. Environmental factors that influence the transition from lysogenic to lytic existence in the phiHSIC/Listonella pelagia marine phage-host system. Microb Ecol 52, 217–225 (2006).

41. Cabello-Yeves, P. J. et al. Novel Synechococcus Genomes Reconstructed from Freshwater Reservoirs. Front Microbiol 8, 1151 (2017).

42. Chénard, C. & Suttle, C. A. Phylogenetic Diversity of Sequences of Cyanophage Photosynthetic Gene psbA in Marine and Freshwaters. Applied and Environmental Microbiology 74, 5317–5324 (2008).

43. Kavagutti, V. S., Andrei, A.-Ş., Mehrshad, M., Salcher, M. M. & Ghai, R. Phage-centric ecological interactions in aquatic ecosystems revealed through ultra-deep metagenomics. Microbiome 7, 135 (2019).

44. Palermo, C. N., Shea, D. W. & Short, S. M. Analysis of Different Size Fractions Provides a More Complete Perspective of Viral Diversity in a Freshwater Embayment. Applied and Environmental Microbiology 87, e00197–21 (2021).

45. Brum, J. R. et al. Ocean plankton. Patterns and ecological drivers of ocean viral communities. Science 348, 1261498 (2015).

46. Genomic analysis of uncultured marine viral communities | PNAS. https://www.pnas.org/doi/full/10.1073/pnas.202488399.

47. Thurber, R. V., Haynes, M., Breitbart, M., Wegley, L. & Rohwer, F. Laboratory procedures to generate viral metagenomes. Nat Protoc 4, 470–483 (2009).

48. Butina, T. V. et al. Extended Evaluation of Viral Diversity in Lake Baikal through Metagenomics. Microorganisms 9, 760 (2021).

49. Gu, C. et al. Saline lakes on the Qinghai-Tibet Plateau harbor unique viral assemblages mediating microbial environmental adaption. iScience 24, 103439 (2021).

50. Potapov, S. et al. The Viral Fraction Metatranscriptomes of Lake Baikal. Microorganisms 10, 1937 (2022).

51. Roux, S. et al. Assessing the Diversity and Specificity of Two Freshwater Viral Communities through Metagenomics. PLoS ONE 7, e33641 (2012).

52. Moon, K., Kim, S., Kang, I. & Cho, J.-C. Viral metagenomes of Lake Soyang, the largest freshwater lake in South Korea. Sci Data 7, 349 (2020).

53. Aguirre de Carcer, D., Lopez-Bueno, A., Pearce, D. A. & Alcami, A. Biodiversity and distribution of polar freshwater DNA viruses. Sci Adv 1, e1400127 (2015).

54. Mohiuddin, M. & Schellhorn, H. E. Spatial and temporal dynamics of virus occurrence in two freshwater lakes captured through metagenomic analysis. Front Microbiol 6, 960 (2015).

55. Kim, Y., Aw, T. G., Teal, T. K. & Rose, J. B. Metagenomic Investigation of Viral Communities in Ballast Water. Environ. Sci. Technol. 49, 8396–8407 (2015).

56. Elbehery, A. H. A. & Deng, L. Insights into the global freshwater virome. Frontiers in Microbiology 13, (2022).

57. Moon, K. et al. Freshwater viral metagenome reveals novel and functional phage-borne antibiotic resistance genes. Microbiome 8, 75 (2020).

58. Chopyk, J. et al. Seasonal dynamics in taxonomy and function within bacterial and viral metagenomic assemblages recovered from a freshwater agricultural pond. Environmental Microbiome 15, 18 (2020).

59. Bibby, K. et al. Metagenomics and the development of viral water quality tools. npj Clean Water 2, 1–13 (2019).

60. Roux, S., Hallam, S. J., Woyke, T. & Sullivan, M. B. Viral dark matter and virus–host interactions resolved from publicly available microbial genomes. eLife 4, e08490 (2015).

61. Berdjeb, L., Ghiglione, J.-F. & Jacquet, S. Bottom-Up versus Top-Down Control of Hypo- and Epilimnion Free-Living Bacterial Community Structures in Two Neighboring Freshwater Lakes. Applied and Environmental Microbiology 77, 3591–3599 (2011).

62. Aeschbach-Hertig, W., Hofer, M., Schmid, M., Kipfer, R. & Imboden, D. M. The physical structure and dynamics of a deep, meromictic crater lake (Lac Pavin, France). Hydrobiologia 487, 111–136 (2002).

63. Jaffe, A. L. et al. Variable impact of geochemical gradients on the functional potential of bacteria, archaea, and phages from the permanently stratified Lac Pavin. 2022.07.18.500538 Preprint at https://doi.org/10.1101/2022.07.18.500538 (2022).

64. Jenny, J.-P. et al. Inherited hypoxia: A new challenge for reoligotrophicated lakes under global warming. Global Biogeochemical Cycles 28, 1413–1423 (2014).

65. Sorek, R., Kunin, V. & Hugenholtz, P. CRISPR--a widespread system that provides acquired resistance against phages in bacteria and archaea. Nat Rev Microbiol 6, 181–186 (2008).

66. Koskella, B. & Meaden, S. Understanding Bacteriophage Specificity in Natural Microbial Communities. Viruses 5, 806–823 (2013).

67. Amundson, K. K., Roux, S., Shelton, J. L. & Wilkins, M. J. Long-term CRISPR array dynamics and stable host-virus co-existence in subsurface fractured shales. http://biorxiv.org/lookup/doi/10.1101/2023.02.03.526977 (2023) doi:10.1101/2023.02.03.526977.

68. Wilhelm, S. W. et al. Marine and Freshwater Cyanophages in a Laurentian Great Lake: Evidence from Infectivity Assays and Molecular Analyses of g20 Genes. Appl Environ Microbiol 72, 4957–4963 (2006).

69. Denef, V. J., Mueller, R. S., Chiang, E., Liebig, J. R. & Vanderploeg, H. A. Chloroflexota CL500-11 Populations That Predominate Deep-Lake Hypolimnion Bacterioplankton Rely on Nitrogen-Rich Dissolved Organic Matter Metabolism and C1 Compound Oxidation. Appl Environ Microbiol 82, 1423–1432 (2015).

70. Mehrshad, M. et al. Hidden in plain sight—highly abundant and diverse planktonic freshwater Chloroflexota. Microbiome 6, 176 (2018).

71. Genomic and Seasonal Variations among Aquatic Phages Infecting the Baltic Sea Gammaproteobacterium Rheinheimera sp. Strain BAL341 | Applied and Environmental Microbiology. https://journals.asm.org/doi/10.1128/AEM.01003-19.

72. Linz, A. M., Aylward, F. O., Bertilsson, S. & McMahon, K. D. Time-series metatranscriptomes reveal conserved patterns between phototrophic and heterotrophic microbes in diverse freshwater systems. Limnology and Oceanography 65, S101–S112 (2020).

73. Kopylova, E., Noé, L. & Touzet, H. SortMeRNA: fast and accurate filtering of ribosomal RNAs in metatranscriptomic data. Bioinformatics 28, 3211–3217 (2012).

74. Haft, D. H. et al. TIGRFAMs and Genome Properties in 2013. Nucleic Acids Res 41, D387–395 (2013).

75. Pruesse, E. et al. SILVA: a comprehensive online resource for quality checked and aligned ribosomal RNA sequence data compatible with ARB. Nucleic Acids Research 35, 7188– 7196 (2007).

76. Nurk, S., Meleshko, D., Korobeynikov, A. & Pevzner, P. A. metaSPAdes: a new versatile metagenomic assembler. Genome Res. 27, 824–834 (2017).

77. Uritskiy, G. V., DiRuggiero, J. & Taylor, J. MetaWRAP—a flexible pipeline for genome-resolved metagenomic data analysis. Microbiome 6, 158 (2018).

78. Kang, D. D., Froula, J., Egan, R. & Wang, Z. MetaBAT, an efficient tool for accurately reconstructing single genomes from complex microbial communities. PeerJ 3, e1165 (2015).

79. Kang, D. D. et al. MetaBAT 2: an adaptive binning algorithm for robust and efficient genome reconstruction from metagenome assemblies. PeerJ 7, e7359 (2019).

80. Wu, Y.-W., Simmons, B. A. & Singer, S. W. MaxBin 2.0: an automated binning algorithm to recover genomes from multiple metagenomic datasets. Bioinformatics 32, 605–607 (2016).

81. Sieber, C. M. K. et al. Recovery of genomes from metagenomes via a dereplication, aggregation and scoring strategy. Nature Microbiology 3, 836–843 (2018).

82. Chaumeil, P.-A., Mussig, A. J., Hugenholtz, P. & Parks, D. H. GTDB-Tk: a toolkit to classify genomes with the Genome Taxonomy Database. Bioinformatics 36, 1925–1927 (2020).

83. Parks, D. H., Imelfort, M., Skennerton, C. T., Hugenholtz, P. & Tyson, G. W. CheckM: assessing the quality of microbial genomes recovered from isolates, single cells, and metagenomes. Genome research 25, 1043–55 (2015).

84. Olm, M. R., Brown, C. T., Brooks, B. & Banfield, J. F. dRep: a tool for fast and accurate genomic comparisons that enables improved genome recovery from metagenomes through de-replication. The ISME Journal 1–5 (2017) doi:10.1038/ismej.2017.126.

85. The Genome Standards Consortium et al. Minimum information about a single amplified genome (MISAG) and a metagenome-assembled genome (MIMAG) of bacteria and archaea. Nature Biotechnology 35, 725–731 (2017).

86. Woodcroft, B. J. CoverM. (2022).

87. Langmead, B. & Salzberg, S. L. Fast gapped-read alignment with Bowtie 2. Nature Methods 9, 357–359 (2012).

88. Li, H. et al. The Sequence Alignment/Map format and SAMtools. Bioinformatics 25, 2078– 2079 (2009).

89. Olm, M. R. et al. inStrain profiles population microdiversity from metagenomic data and sensitively detects shared microbial strains. Nat Biotechnol 39, 727–736 (2021).

90. Zhou, Z. et al. METABOLIC: high-throughput profiling of microbial genomes for functional traits, metabolism, biogeochemistry, and community-scale functional networks. Microbiome 10, 33 (2022).

91. Kanehisa, M., Furumichi, M., Tanabe, M., Sato, Y. & Morishima, K. KEGG: new perspectives on genomes, pathways, diseases and drugs. Nucleic Acids Research 45, D353– D361 (2017).

92. Mistry, J. et al. Pfam: The protein families database in 2021. Nucleic Acids Research 49, D412–D419 (2021).

93. Yin, Y. et al. DbCAN: A web resource for automated carbohydrate-active enzyme annotation. Nucleic Acids Research 40, 445–451 (2012).

94. Rawlings, N. D. et al. The MEROPS database of proteolytic enzymes, their substrates and inhibitors in 2017 and a comparison with peptidases in the PANTHER database. Nucleic Acids Research 46, D624–D632 (2018).

95. McDaniel, E. A., Anantharaman, K. & McMahon, K. D. metabolisHMM: Phylogenomic analysis for exploration of microbial phylogenies and metabolic pathways. 2019.12.20.884627 Preprint at https://doi.org/10.1101/2019.12.20.884627 (2020).

96. Liu, K., Linder, C. R. & Warnow, T. RAxML and FastTree: Comparing Two Methods for Large-Scale Maximum Likelihood Phylogeny Estimation. PLoS ONE 6, e27731 (2011).

97. Kieft, K., Zhou, Z. & Anantharaman, K. VIBRANT: Automated recovery, annotation and curation of microbial viruses, and evaluation of virome function from genomic sequences. http://biorxiv.org/lookup/doi/10.1101/855387 (2019) doi:10.1101/855387.

98. Nayfach, S. et al. CheckV assesses the quality and completeness of metagenome-assembled viral genomes. Nat Biotechnol 39, 578–585 (2021).

99. Buchfink, B., Xie, C. & Huson, D. H. Fast and sensitive protein alignment using DIAMOND. Nature methods 12, 59–60 (2015).

100. Edgar, R. C. PILER-CR: Fast and accurate identification of CRISPR repeats. BMC Bioinformatics 8, 18 (2007).

