## Supplementary Figures 1-7 for "Viral impacts on microbial activity and biogeochemical cycling in a seasonally anoxic freshwater lake"

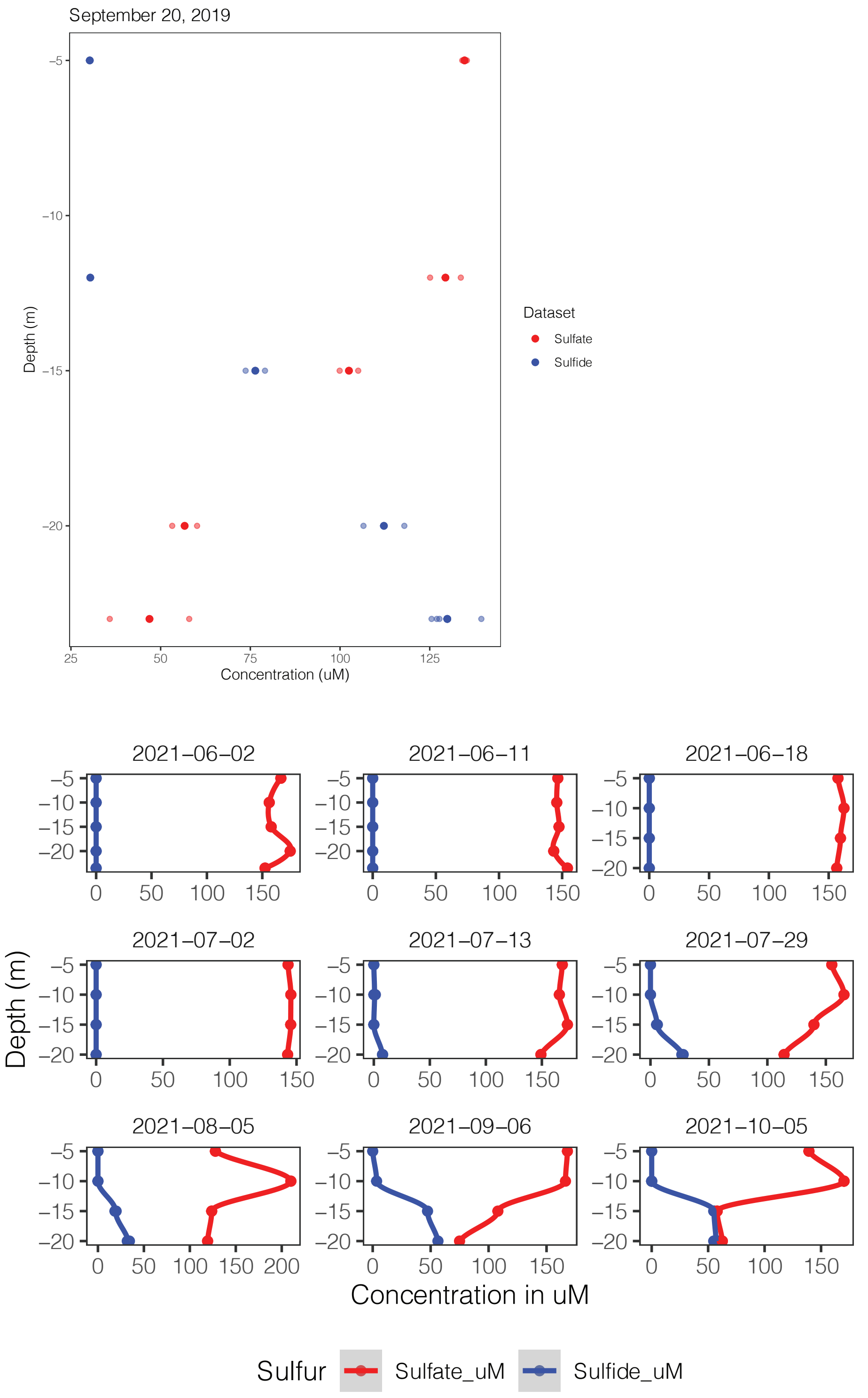


**Supplementary Figure 1**. Sulfate and sulfide profiles (uM) in Lake Mendota in 2019 and 2021.

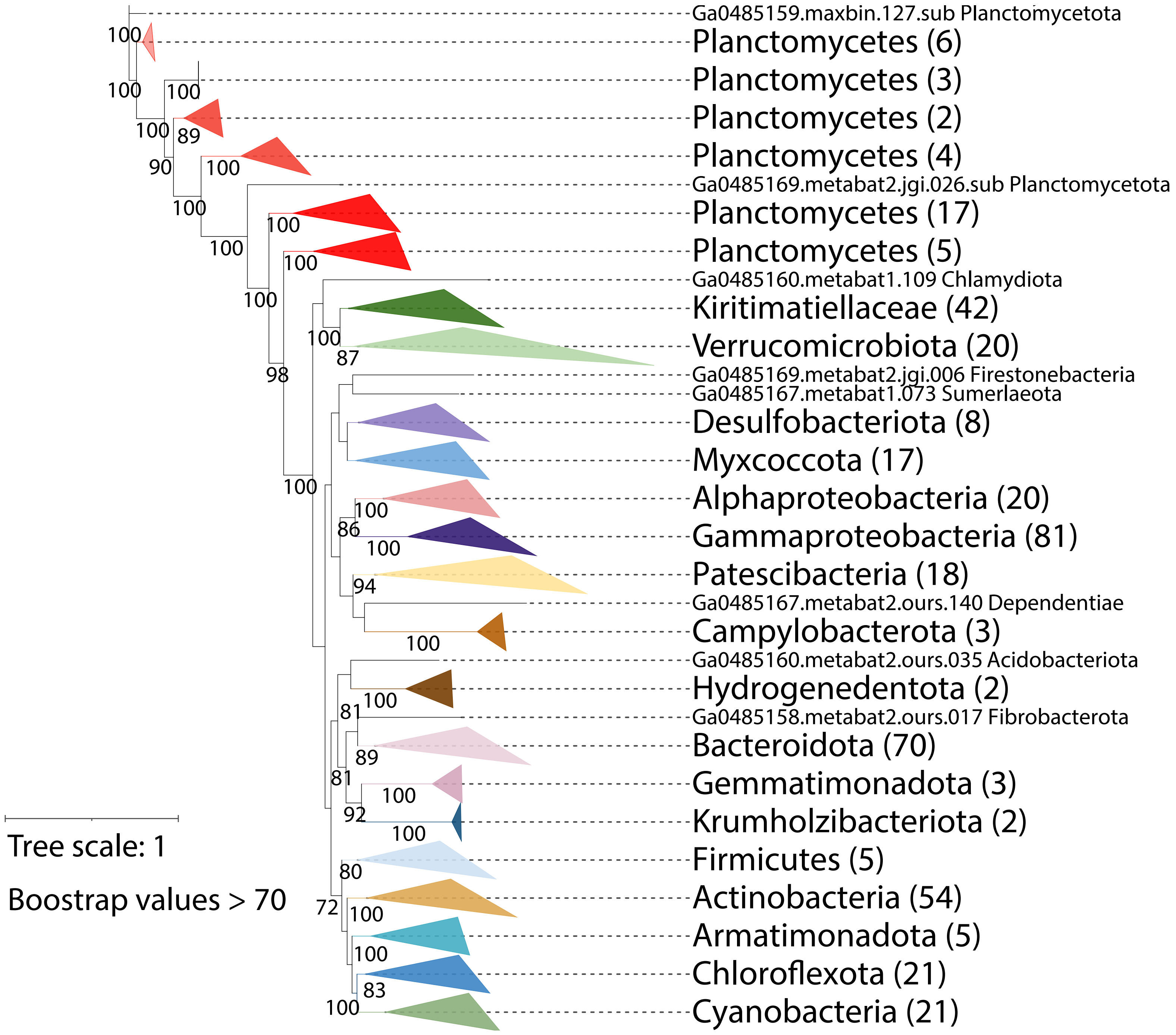


**Supplementary Figure 2. Phylogenetic tree of the 429 Bacterial MAGs using 16 ribosomal proteins. The 2 Archaea are not shown.**

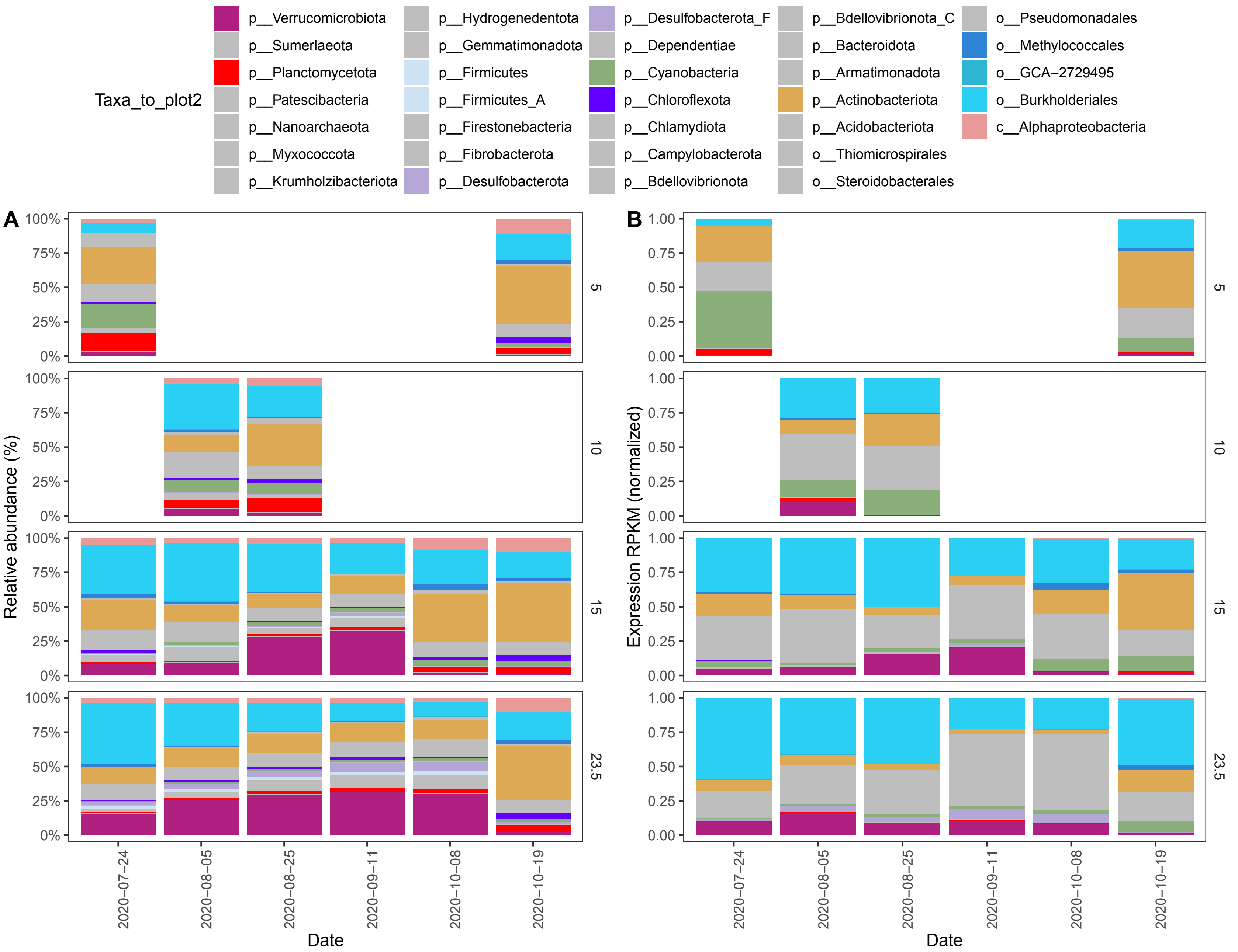


**Supplementary Figure 3. Bar plot showing the abundance and expression of all MAGs in Lake Mendota in this study.**

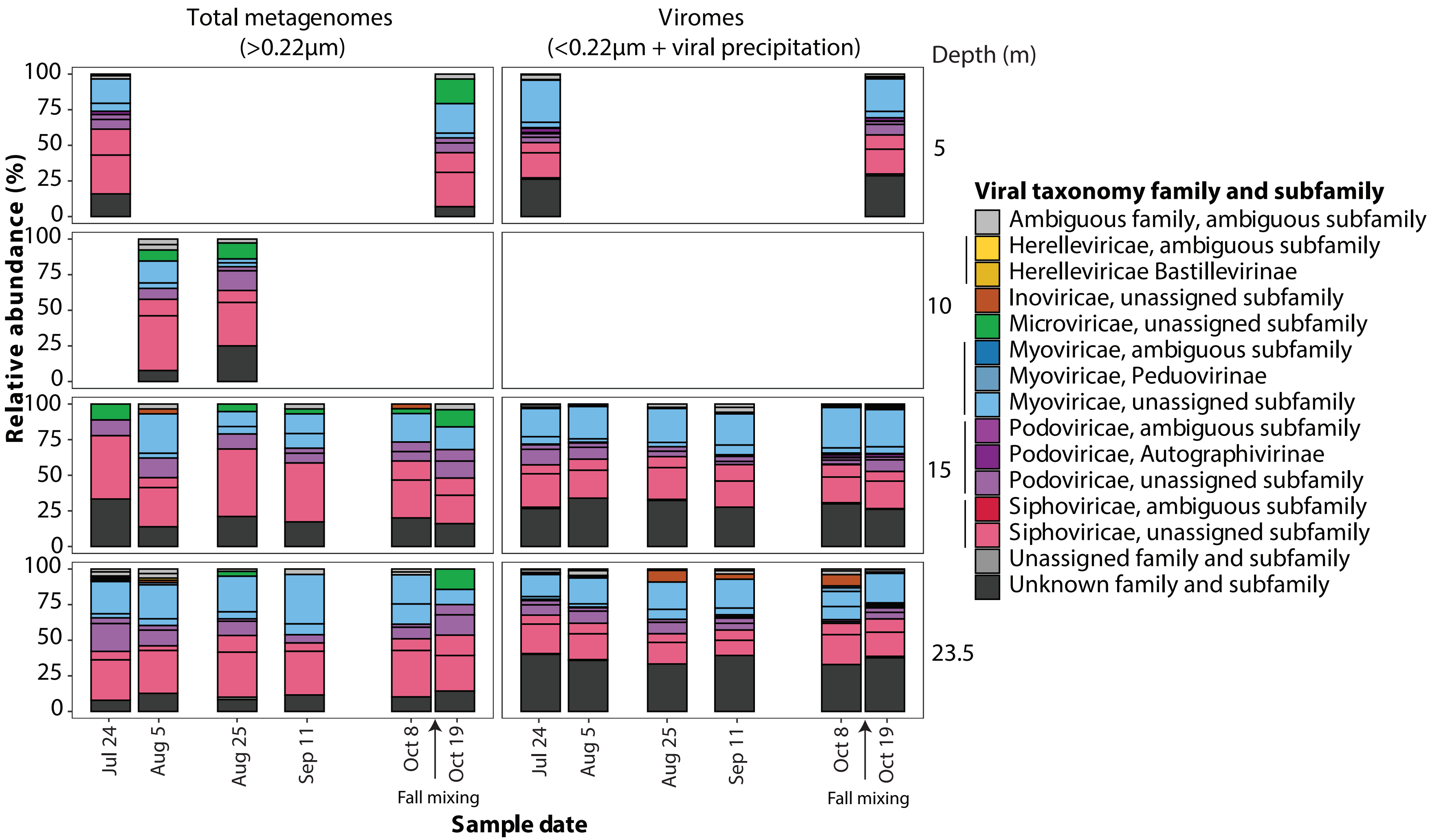


**Supplementary Figure 4**. Bar plot showing the taxonomic composition of the phages in Lake Mendota in this study.

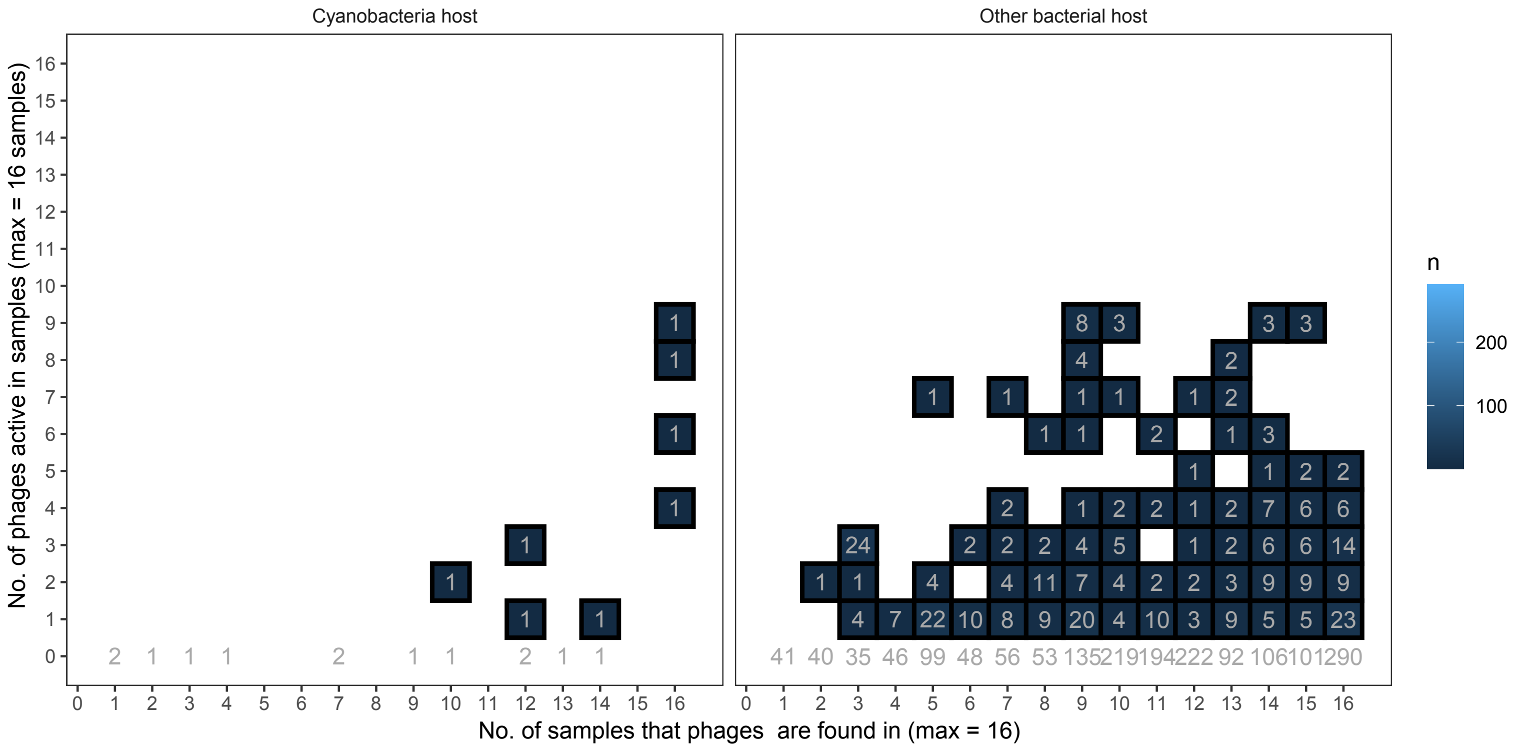


**Supplementary Figure 5**. Number of phages and samples they are expressed or present in.

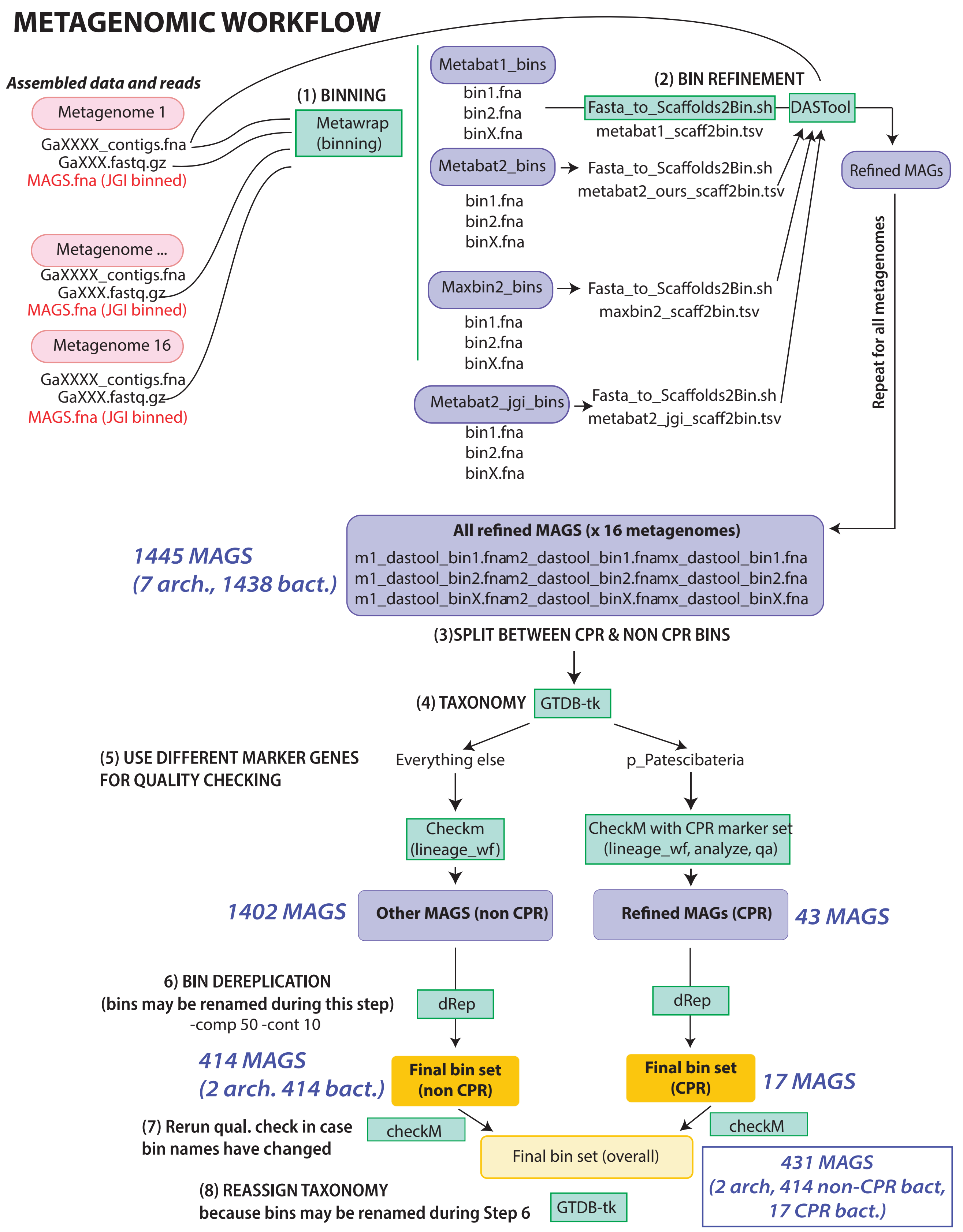


**Supplementary Figure 6**. Metagenomic processing workflow.

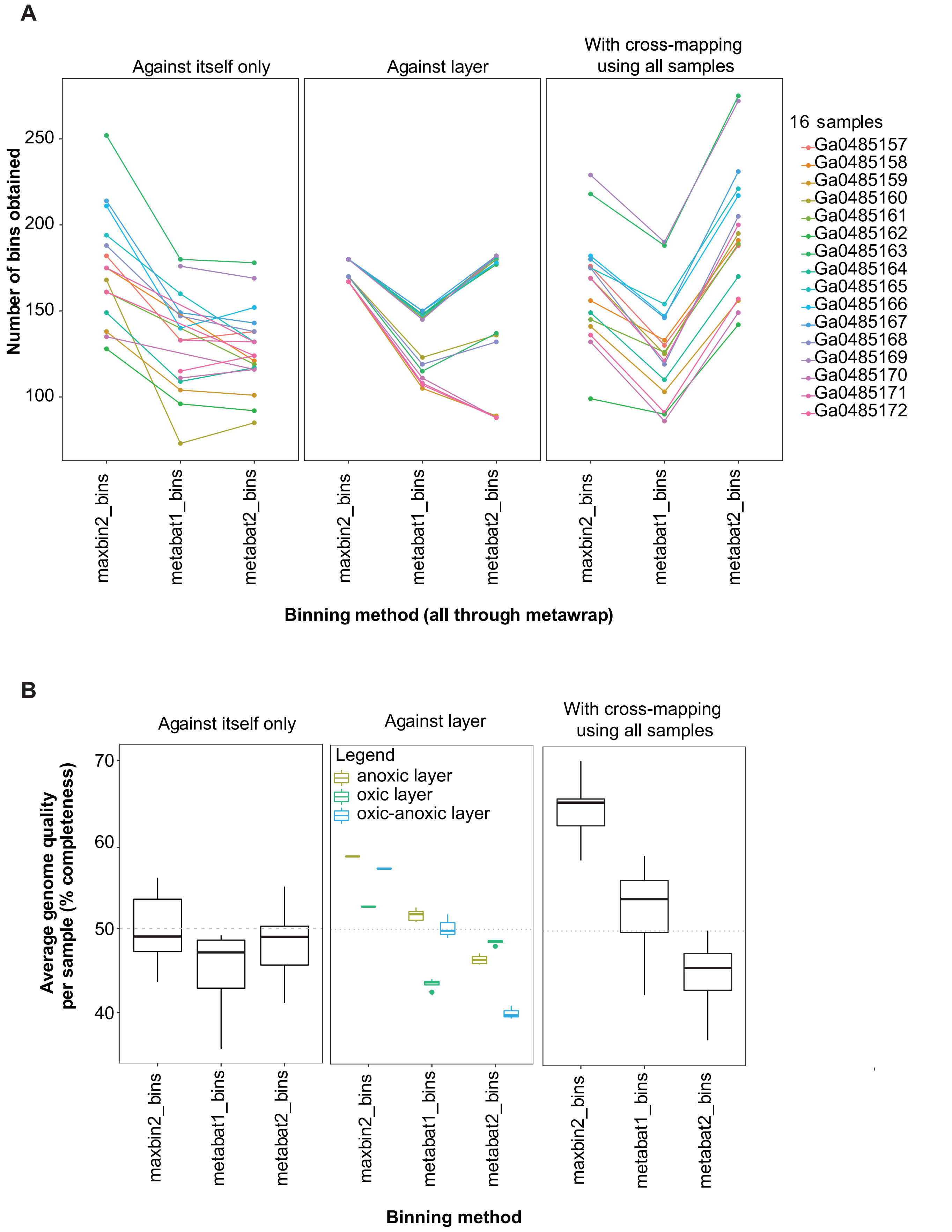


**Supplementary Figure 7**. Comparison of binning using individual sample reads only, reads from samples with similar environmental parameters (e.g. oxygen levels), or differential coverage using all samples. Metabat2 with cross-mapping resulted in an overall higher number of MAGs (A), but maxbin2 with cross-mapping resulted in the highest overall quality of MAGs (B). For this reason, cross-mapping was chosen as the method to obtain more, and high-quality MAGs, before refinement and dereplication.
